# Enhancing Myocardial Repair with CardioClusters

**DOI:** 10.1101/759845

**Authors:** Megan M. Monsanto, Bingyan J. Wang, Zach R. Ehrenberg, Oscar Echeagaray, Kevin S. White, Roberto Alvarez, Kristina Fisher, Sharon Sengphanith, Alvin Muliono, Natalie A. Gude, Mark A. Sussman

## Abstract

**Background:** Cellular therapy to treat heart failure is an ongoing focus of intense research and development, but progress has been frustratingly slow due to limitations of current approaches. Engineered augmentation of established cellular effectors overcomes impediments, enhancing reparative activity with improved outcomes relative to conventional techniques. Such ‘next generation’ implementation includes delivery of combinatorial cell populations exerting synergistic effects. Concurrent isolation and expansion of three distinct cardiac-derived interstitial cell types from human heart tissue, as previously reported by our group, prompted design of a three-dimensional (3D) structure that maximizes cellular interaction, allows for defined cell ratios, controls size, enables injectability, and minimizes cell losses upon delivery.

**Methods:** Three distinct populations of human cardiac interstitial cells including mesenchymal stem cells (MSCs), endothelial progenitor cells (EPCs), and c-Kit^+^ cardiac interstitial cells (cCICs) when cultured together spontaneously form scaffold-free 3D microenvironments termed CardioClusters. Biological consequences of CardioCluster formation were assessed by multiple assays including single cells RNA-Seq transcriptional profiling. Protective effects of CardioClusters in vitro were measured using cell culture models for oxidative stress and myocardial ischemia in combination with freshly isolated neonatal rat ventricular myocytes. Long-term impact of adoptively transferred CardioClusters upon myocardial structure and function in a xenogenic model of acute infarction using NOD^scid^ mice was assessed over a longitudinal time course of 20-weeks.

**Results:** CardioCluster design enables control over composite cell types, cell ratios, size, and preservation of structural integrity during delivery. Profound changes for biological properties of CardioClusters relative to constituent parental cell populations include enhanced expression of stem cell-relevant factors, adhesion/extracellular-matrix molecules, and cytokines. The CardioCluster 3D microenvironment maximizes cellular interaction while maintaining a more native transcriptome similar to endogenous cardiac cells. CardioCluster delivery improves cell retention following intramyocardial injection with preservation of long-term cardiac function relative to monolayer-cultured cells when tested in an experimental murine infarction model followed for up to 20 weeks post-challenge. CardioCluster-treated hearts show increases in capillary density, preservation of cardiomyocyte size, and reduced scar size indicative of blunting pathologic infarction injury.

**Conclusions:** CardioClusters are a novel ‘next generation’ development and delivery approach for cellular therapeutics that potentiate beneficial activity and enhance protective effects of human cardiac interstitial cell mixed populations. CardioClusters utilization in this preclinical setting establishes fundamental methodologic and biologic insights, laying the framework for optimization of CardioCluster design to provide greater efficacy in cell-based therapeutic interventions intended to mitigate cardiomyopathic damage.

## Introduction

Cellular therapy continues to be pursued as an experimental approach to treat cardiomyopathic damage following infarction in both preclinical animal models and human clinical trials^1–9^. While promising, cellular therapy has been hindered by marginal improvement in cardiac function, in part due to low engraftment and persistence of the transplanted cells^10–12^. As such, improving cell retention and survival has been a major area of research using a plethora of techniques, including biomaterials^13, 14^, cytokines and growth factors^15^, repeated administration of cells^16^, or genetic enhancement with pro-survival and anti-apoptotic genes^17, 18^. Combinatorial cell therapies are an additional conceptual approach intended to promote additive or synergistic effects in myocardial repair^19–22^; however, cell retention and survival remain poor whether single or combinatorial cell delivery is attempted. Furthermore, coincident delivery of multiple cell populations does not ensure effective cell interaction and cross talk following injection. Therefore, novel strategic approaches to prolong cell retention and survival are highly sought after, preferably also favoring cell phenotypic characteristics that promote short-term mitigation of injury and long-term recovery of myocardial structure and function.

Traditional monolayer culture, while both prevalent and convenient, promotes loss of cell identity and spatial organization, thereby impacting many cellular aspects, including cell morphology, proliferation, differentiation, viability, and transcriptome profile^23–25^. Approaches intended to mitigate deleterious effects of artificial *in vitro* environments through recreating native three dimensional (3D) architecture include engineered heart tissue grafts^26, 27^, organoids^28, 29^, and 3D bioprinting.^30, 31^ When grown in 3D cells aggregate and form complex cell-cell and cell-matrix interactions that mimic the natural environment found *in vivo,* allowing for cell differentiation and tissue organization not possible in conventional two-dimensional (2D) culture systems^32^. For example, cardiosphere formation^33, 34^ enhances cellular communication through gap junction protein connexin 43 fostering signal propagation and cell differentiation^35–37^. However, cardiospheres are formed by random cell aggregation leading to inconsistent cell characterization markers^33, 34^, variable sphere size, and muddled communication within the aggregate microenvironment^33, 34, 38^. A superior preferable methodology would utilize combinations of defined cell populations aggregated in 3D under controlled conditions that allow for intact delivery without dissociation of the spherical microenvironment to promote preservation of cellular phenotype and optimize cellular retention. This combination of desired features formed the rational basis for our novel methodological approach distinct in several critical aspects from cardiosphere-derived cell therapy.

A ‘next generation’ conceptual approach for a deliverable cell therapeutic based upon the aforementioned literature precedents would involve 1) combinations of multiple cardiac-derived cell types, 2) *ex vivo* culture adaptation to a 3D microenvironment to maximize cellular interaction, 3) control over cell composition, cell number, and aggregate cell volume, and 4) capability to deliver *ex vivo* formed cell aggregate ‘niches’ intact to facilitate preservation of microenvironment biological properties and retention. Guided by these stipulated parameters to create a novel artificially engineered cell aggregate, inception began with essential methodological studies to isolate and expand three distinct cardiac-derived interstitial cell types from a single human heart biopsy as previously reported by our group^39^. Human cardiac tissue of varying age and gender was used as source material to isolate c-Kit^+^ cardiac interstitial cells (cCIC), mesenchymal stem cells (MSCs), and endothelial progenitor cells (EPCs). Having isolated and characterized three distinct cardiac interstitial cell types each known to possess beneficial properties to blunt cardiomyopathic damage, a protocol was designed for spontaneous self-assembly of mixed cell populations into an optimal conceptual engineered solution that we have named “CardioClusters.”

CardioClusters harness the distinct phenotypic attributes of three well defined cardiac cell populations. MSCs support myocardial reparative activity through secretion of paracrine factors that activate endogenous cells, promote angiogenesis, protect cardiomyocytes, and reduce scar formation^40, 41^. MSCs also secrete cell adhesion molecules such as integrins and cadherins integral to cellular aggregation^42^. EPCs promote paracrine dependent vasculogenesis and angiogenesis and differentiate into mature endothelial cells^43, 44^. Prior studies demonstrate EPCs transplanted *in vivo* are capable of forming microvessels, but regress without MSCs to support vessel maturity^45, 46^. cCICs distributed in the CardioCluster contribute to support of myocardial homeostasis, response to injury, and remodeling. CardioCluster characteristics are particularly well suited to mediating myocardial repair in the wake of acute pathological damage by providing a more natural niche microenvironment for augmented delivery and extended functional activity to improve the outcome of cellular therapy.

CardioClusters advance the application of combinatorial cell therapy by integrating complementary and synergistic properties from multiple cardiac-resident cell types into a single injectable product. CardioCluster formation is a rapid, reproducible, and controllable process demonstrated in preclinical testing to mediate significant improvements in myocardial structure and function following infarction injury. Enhancement of cell therapy with CardioClusters represents an important advance to improve clinical application of cell therapy for treatment of cardiomyopathic damage and disease.

## Methods

Full materials and methods are available in the Online Data Supplements.

### Human Cardiac Interstitial Cell Isolation

NIH guidelines for human research were followed as approved by IRB review (Protocol #120686). Neonatal heart tissue procured from post-mortem infants provided by a commercial source (Novogenix Laboratories) was used for isolation of human cardiac cells. The protocol detailing cardiac cell isolation can be found in our previous publication^39^ and Online Data Supplement. All cells used in this study were mid-passage (passages 5–10).

### CardioCluster Formation

CardioClusters are formed using 96 well, ultra-low attachment multiwell round-bottom plates (Corning, catalog #CLS7007) in a two-step process. Step 1 generates the inner core composed of cCICs and MSCs in a 1:2 ratio. The inner core of cCICs and MSCs is seeded in 100 µL/well MSC media for 24 hours at 37°C in 5% CO_2_ incubator. Step 2 forms the outer EPC layer using a cell number equal to the number of cells used to create the central core. The EPCs are added in an additional 50 µL/well MSC media and incubated at 37°C in 5% CO_2_ for an additional 24 hours until CardioCluster 3D structure has formed. The radius of a CardioCluster is approximately 150 μm, composed of a total of 400±100 cells.

### Single-cell RNA-seq Preparation, Data Analysis, and Data Availability

Size distribution was quantified for single cell suspensions of cCIC, MSC, EPC and dissociated CardioClusters to verify cell size met droplet platform specifications (Supplemental Figure 6A). Cells were loaded on a Chromium™ Controller (10x Genomics) and single-cell RNA-Seq libraries were prepared using Chromium™ Single Cell 3’ Library & Gel Bead Kit v2 (10x Genomics) following manufacturer’s protocol. Each library was tested with Bioanalyzer (average library size: 450-490 bp). The sequencing libraries were quantified by quantitative PCR (KAPA Biosystems Library Quantification Kit for Illumina platforms P/N KK4824) and Qubit 3.0 with dsDNA HS Assay Kit (Thermo Fisher Scientific). Sequencing libraries were loaded at 2 pM on an Illumina HiSeq2500 with 2X75 paired-end kits using the following read length: 98 bp Read1, 8 bp i7 Index, and 26 bp Read2. Batch effects concerns were obviated in our analyses by mitigating strategies in our experimental design. Specifically, the following steps were taken: 1) all samples were processed in the same microfluidics chip, and 2) libraries were prepared in parallel and sequenced in the same lane in the HiSeq 4000. Libraries were aggregated and normalized using CellRanger. Final removal of unwanted sources of variation and batch effect corrections was performed using Seurat R Package (v2.3.4)^47^.

Raw sequencing data was processed with the Cell Ranger pipeline (10X Genomics; version 2.0). Sequencing reads were aligned to the human genome hg19. Cells with fewer than 1,000 genes or more than 10% of mitochondrial gene UMI count were filtered out and genes detected in fewer than three cells were filtered out using Seurat R Package (v2.3.4)^47^. The first 20 principal components were found to be significant to perform dimensionality reduction. Preparations derived from *in vitro* studies yielded 5659 barcoded cells for analysis, from which 1125, 1717, 1403 and 1414 corresponded to CardioClusters, cCICs, EPCs and MSCs, respectively. Approximately 2,029 variable genes were selected based on their expression and dispersion. The first 20 principal components were used for the t-SNE projection and unsupervised clustering^47^. Differential expression analysis was done using Wilcoxon rank sum test and selecting for an adjusted p-value ≤0.05 and a log (FC)>0.25. Global differential expression analysis was done using Loupe Cell Browser 2.0.0. Gene ontology analysis was performed using R package clusterProfiler^48^. scRNA-Seq data generated in this study have been uploaded to the Gene Expression Omnibus (GEO) database (accession number GSE133832). Datasets for freshly isolated CICs have been previously published by our group^25^ and are available at the GEO database (accession number GSE114280).

### Myocardial Infarction and Intramyocardial Injection

Animal protocols and experimental procedures are approved by the Institutional Animal Care and Use Committee at San Diego State University. Animals were randomized for treatments that were blinded to personnel carrying out the surgical procedures, injections, and physiological function analysis. Appropriate animal sample size was determined using sample size calculator (http://www.lasec.cuhk.edu.hk/sample-size-calculation.html). A total of 60 mice were used in this study. Inclusion criteria were set prior to study commencement and required a drop in ejection fraction between 10-50% 1-week following infarction injury. Myocardial infarctions were carried out on 8-week old NOD.CB17-Prkdc^scid^/J female mice (The Jackson Laboratory, catalog #001303) under 2% isoflurane (Victor Medical, catalog #NDC 57319-474-06) as previously described^49^. Briefly, the 3rd and 4th ribs were separated enough to get adequate exposure of the operating region, but the ribs were kept intact. The heart was squeezed out by pressing the thorax lightly and the left anterior descending artery (LAD) was ligated at the distal diagonal branch with a 7-0 suture. Infarction was confirmed by blanching of anterior left myocardium wall. Following ligation, either CardioClusters (n=17), 2:3:1 ratio of cCIC+EPC+MSC (C+E+M; n=15), or a vehicle control (PBS plus 0.5% sodium alginate [NovaMatrix, catalog #4209001]; n=16) were delivered intramyocardially at 3 separate sites in the vicinity bordering the blanched area. A total of 90,000 cells per heart were introduced into CardioCluster and C+E+M animals. The hearts were immediately placed back into the intrathoracic space followed by muscle and skin closure. Animals in sham group (n=12) received a comparable surgical procedure without LAD ligation or injection. Each animal received 20 µL analgesic treatment with 0.3 mg/mL Buprenex (Victor Medical, catalog #12496-0757) at time of surgery and 12 hours post-surgery. No animals required exclusion from study.

### Statistical Analysis

Data are expressed as mean±SEM. Statistical analyses of multiple groups were assessed by 1-way ANOVA with Bonferroni post hoc test (comparison among all groups) or Dunnett’s post hoc test (versus single group). Multiple groups over time were analyzed by 2-way ANOVA. Statistical analysis was performed using GraphPad Prism version 5.0 software. Experiments were performed in triplicate unless stated otherwise. A p-value of less than 0.05 was considered statistically significant.

## Results

### Three distinct cardiac nonmyocyte cell types are used for CardioCluster formation

CardioClusters are comprised of three cardiac-resident cell populations concurrently isolated as previously described by our group with full phenotypic characterization^39^. Cell surface marker profiling of cell lines used to produce CardioClusters were similar to previous findings (cCIC express c-Kit^high^, CD90^high^, CD105^low^, CD133^low^, CD45^neg^; EPC express CD133^high^, CD105^high^, c-Kit^low^, CD90^neg^, CD45^neg^; and MSC express CD90^high^, CD105^high^, c-Kit^low^, CD133^low^, CD45^neg^; data not shown). Commitment toward angiogenic and smooth muscle fates was assessed for the three cardiac-derived cell populations relative to control cell lines human umbilical vein endothelial cells (HUVECs) and bone marrow-derived MSCs (BM MSCs). Tube formation assays demonstrated robust angiogenic responses from both EPCs and HUVECs *in vitro* using growth factor-reduced Matrigel (Supplemental Figure 1A). Transcript levels of endothelial-related genes *CD31* and *von Willebrand factor* (*vWF*) were elevated in HUVECs and EPCs (p<0.001 versus cCICs; Supplemental Figure 1B and 3C). *Smooth muscle actin* (*SMA*) transcripts were highly expressed in BM MSCs and cardiac MSCs (p<0.001 versus cCICs), with both endothelial populations (EPC or HUVEC) expressing near undetectable levels (p<0.05 versus cCICs; Supplemental Figure 1D). *GATA4* was expressed by cCICs (1.0±0.05) and to a lesser extent by EPCs (0.87±0.03) and MSCs (0.33±0.01), with non-cardiac controls expressing undetectable levels (Supplemental Figure 1E). Collectively, these three cardiac-derived cell populations recapitulate and validate previous results of phenotypic characterization for cell types obtained using our published protocol^39^. Distinct phenotypic properties of these three cardiac-derived cell populations fulfills the conceptual design of combining multiple cell types for CardioClusters formation.

The three distinct cardiac derived cell populations were modified with lentiviral vectors to introduce fluorescent proteins for tracking purposes (eGFP tagged cCICs [green], mOrange tagged EPCs [blue], and Neptune tagged MSCs [red]; tagging efficiency 99.1±0.2%; Supplemental Figure 2A and 2B). Distinct morphology for each cell population is evident in representative brightfield images with companion immunofluorescent images demonstrating corresponding fluorophore expression in cCICs (Figure 1A), EPCs (Figure 1B), and MSCs (Figure 1C). Cell morphology measurement of area, roundness, and L/W ratio for each cell type confirmed distinct phenotypes (Figure 1, D-F). MSCs were significantly larger (18,563±1,021) relative to both cCIC (3383±121) and EPC (3272±102) (Figure 1D). EPCs were significantly rounder (EPC, 0.55±0.012; cCIC, 0.19±0.0097; MSC, 0.36±0.015) (Figure 1E), while cCICs show increased L/W ratio (cCIC, 5.2±0.19; EPC, 2.1±0.063; MSC, 2.8±0.11) (Figure 1F). Morphometric parameters clustered by cell type (Supplemental Figure 3), with minor variation between heart samples. EPCs exhibited a proliferative rate similar to cCICs, with both populations showing increased proliferation over MSCs based on CyQuant proliferation assays (Figure 1G). EPCs were significantly more resistant to cell death and retained 92±0.76% cell viability, versus only 54±5.6% for cCIC and 79±1.5% for MSCs after 4 hours H_2_O_2_ treatment (Figure 1, H-J). Cumulatively, characterization showed phenotypic and biological distinctions between cardiac interstitial cell populations fundamental to CardioCluster design and utility, such as elevated resistance to oxidative stress-induced cell death, high proliferative activity, and pro-angiogenic nature of EPCs.

**Figure 1.**
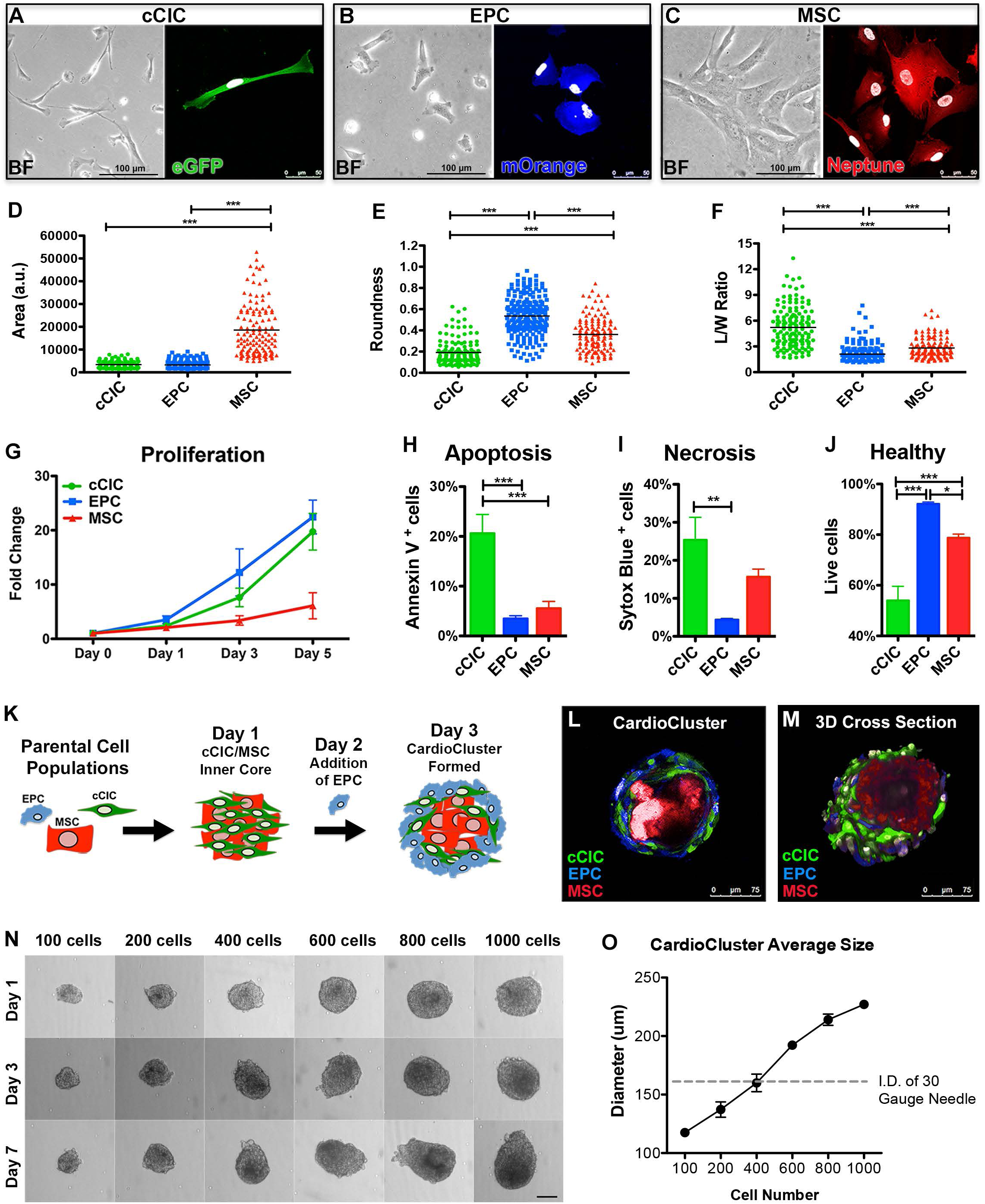
Three distinct cardiac cell types used to generate CardioClusters. **A-C,** Representative brightfield (BF) and immunofluorescent images for cCIC (eGFP^+^) **(A),** EPC (mOrange^+^) **(B)** and MSC (Neptune^+^) **(C).** Scale bars: brightfield, 100 µm; immunofluorescent, 50 µm. DAPI to visualize nuclei (white). **D-F,** Cell morphometric parameters measuring area (a.u.: arbitrary units; **D),** roundness **(E),** and length-to-width (L/W) ratio **(F;** n=3 human heart isolations; minimum of 30 cells traced per cell type, per heart). **G,** Cardiac cell proliferation measured using CyQuant assays at day 0, day 1, day 3, and day 5 (n=3). **H-J,** Annexin V/Sytox Blue labeling for apoptotic **(H),** necrotic **(I)** and healthy **(J)** cells following cell death assay: 24 hours of low serum (75% serum reduction) followed by 4 hours treatment with 30 µM H_2_O_2_ in low serum medium (n=4). Data are presented as 1-way ANOVA, *p<0.05, **p<0.01, ***p<0.001. **K,** Schematic showing CardioCluster formation using three cell populations isolated from human heart tissue: MSC (red), cCIC (green), EPC (blue). **L,** Live CardioCluster visualized by endogenous fluorescent tags showing cCIC (eGFP; FITC channel; green), MSC (Neptune; APC channel; red) and EPC (mOrange; PE channel; blue). Scale bar, 75 µm. **M,** 30 cross section of a CardioCluster lightly fixed in 4% paraformaldehyde and stained with 4’,6-diamidino-2-phenylindole (DAPI; white) to visualize nuclei and cells exhibiting fluorescent tags without the need for antibody labeling. Scale bar, 75 µm. **N,** Representative brightfield images of CardioClusters ranging in size from 100-1000 cells cultured over a 7-day period. Scale bar, 100 µm. **O,** Determination of cell number enabling CardioClusters to pass through a 30-gauge needle, which has an inner diameter (I.D.) of 159 µm. Graph plot of CardioCluster diameters averaged over 3 days (n=4-7).

### Generation of CardioClusters

CardioClusters are formed in a two-step process (Figure 1K). cCICs and MSCs are seeded to form the inner core, with EPCs added 24 hours later to provide an endothelial cell-enriched outer layer for the CardioCluster. The outer EPC layer provides enhanced resistance to oxidative stress relative to the more sensitive cCICs and MSCs within the CardioCluster core (Figure 1J). Individual cells within an assembled CardioCluster were readily visualized with their cognate fluorophore tags, obviating the need for antibody-mediated detection (Figure 1L and 1M). CardioCluster size reproducibly and predictably corresponds to cell number seeded per microwell (Figure 1N). CardioClusters ranging from 100-1000 cells were examined to determine changes in size and morphology over a 7 day time period (Supplemental Figure 2D). CardioCluster diameter and area increased over 7 days, except for CardioClusters seeded with 1000 cells, whose diameter plateaued after day 3. This finding, also previously observed with 3D aggregated cells^50^, is consistent with reduced oxygen and nutrient diffusion within dense cellular structures >200 μm^51, 52^.

Preserving CardioCluster 3D structural integrity for intramyocardial delivery to promote intercellular contact and enhance retention is essential to improve upon typical approaches involving dissociated single cell suspensions. Intramyocardial injection for cell delivery in murine hearts uses a standard 30-gauge needle with a 159 μm internal diameter ([I.D.]), so CardioClusters were engineered for a diameter allowing for injection to preserve 3D structure. The maximum number of cells that could comprise a CardioCluster and pass through a 30-gauge needle is 400 cells based on morphometric quantitative analysis (Figure 1O). EPC, cCICs and MSCs were combined in a 3:2:1 ratio, based upon the consideration of larger MSC size occupying relatively more volume relative to cCIC or EPC (Figure 1D). CardioCluster spontaneous self-assembly as revealed using time-lapse video microscopy shows the MSC population immediately migrating towards the central core (Supplemental Video 1). cCIC/MSC interaction was allowed to progress for 24 hours, at which point EPCs were added to interact with established cCIC/MSC cores. Interestingly, EPCs initially form their own clusters instead of adhering to cCIC/MSC cores and then subsequently envelop the cCIC/MSC core (Supplemental video 1 and Supplemental Figure 2). Architecture of an MSC-enriched core was invariant regardless of seeding sequence, as plating of cCIC+EPCs prior to adding MSCs consistently resulted in MSCs migrating and localizing within the CardioCluster core rather than surface (Supplemental Video 2) consistent with the preferential localization of MSC to hypoxic environments.^53, 54^. CardioCluster formation consistently occurs with MSCs in the core and cCIC/EPCs on the outer layers.

CardioClusters possess a high percentage of live cells maintained within the 3D structure (93.9-98% of cells alive; Supplemental Figure 4). Robust vitality of CardioClusters was confirmed by recovery from long term liquid nitrogen storage, where the percentage of live cells was comparable to that of control non-frozen CardioClusters (Supplemental Figure 4A and 4B). When cultured on standard tissue culture-treated plastic, cells adhered and migrated out from the CardioCluster whether frozen or not non-frozen with comparable cell morphology (Supplemental Figure 4C). These findings support “off-the-shelf” feasibility of using frozen banked CardioClusters for therapeutic purposes rather than necessitating *de novo* creation prior to use.

### CardioCluster cells undergo reprogramming toward the transcriptome profile of freshly isolated cardiac interstitial cells

Transcriptional profiling of CardioClusters and their monolayer cultured parental counterparts reveals significant reprogramming consequential to 3D aggregation. Since CardioClusters are heterogeneous cell populations by design, single-cell RNA sequencing (scRNA-seq) was employed to reveal cellular transcriptome heterogeneity within the CardioCluster (Supplemental Figure 5) at the single cell level with a level of resolution not achievable with bulk population analysis. Quality control testing validated parameters of cell size distribution, sequence alignment and filtering of multiplets and dying cells (Supplemental Figure 6). Dimensionality reduction by t-SNE reveals segregation of CardioClusters (orange cluster) distinguished by a unique transcriptome profile separating them from their constituent parental populations, which form their own clusters (red, green, blue clusters representing MSCs, cCICs and EPCs respectively; Figure 2A). Differentially expressed genes (DEGs) are increased in the CardioCluster environment (620) relative to cCIC (296), EPC (167), or MSC (211) (Figure 2B, Supplemental Table 4). Taken together, these results demonstrate CardioClusters distinguishing themselves as a transcriptionally unique population diverged from parental cell lines.

**Figure 2.**
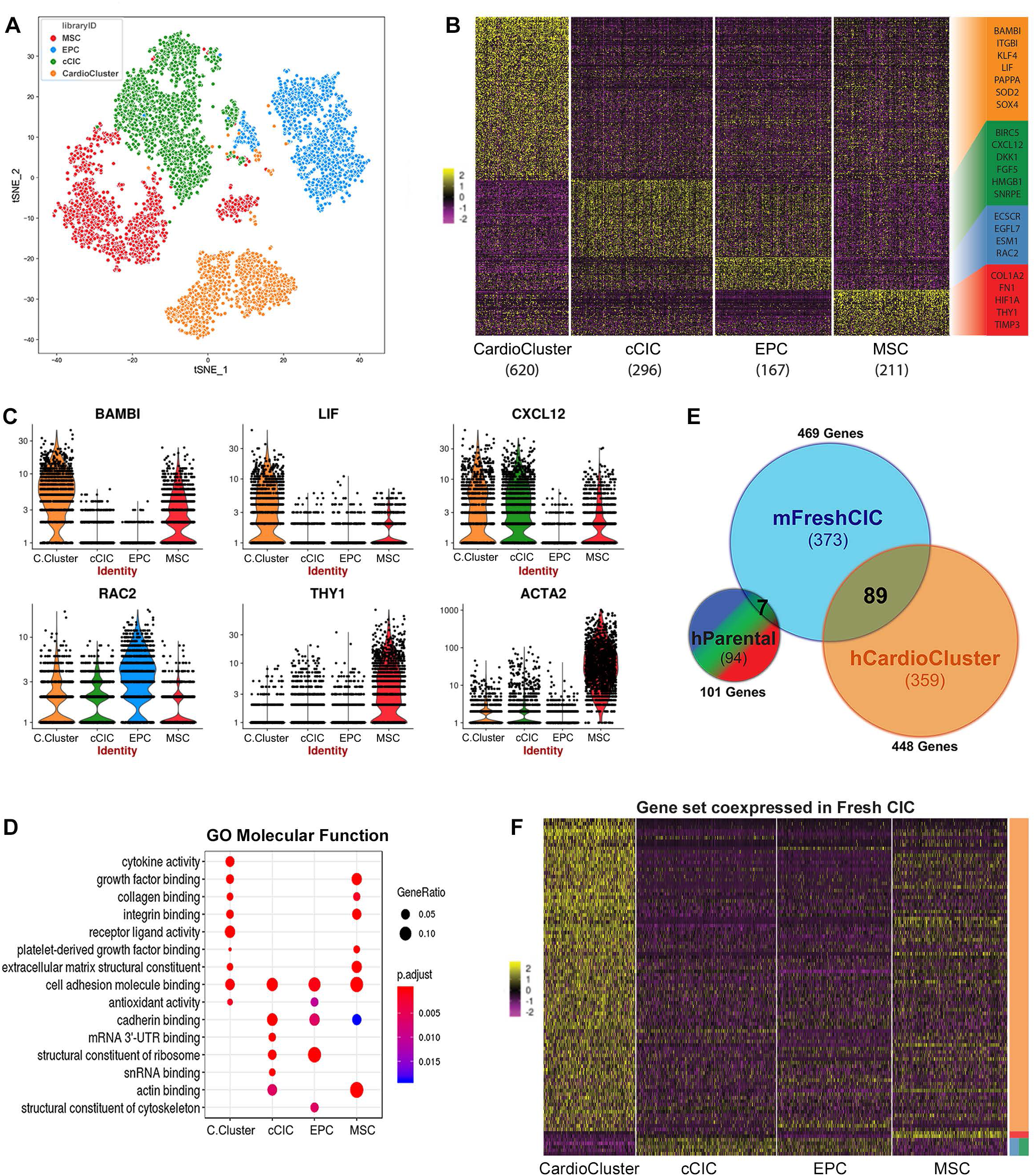
CardioCluster single cell RNA-Seq reveals transcriptional profile acquisition with increased similarity to freshly isolated cardiac interstitial cells. **A-F,** Transcriptional profiling on CardioClusters and parental cells using scRNA-seq. **A,** t-SNE map showing cells grown within a 3D CardioCluster (orange) predominately cluster together, while cCICs (green), EPCs (blue), and MSCs (red) primarily cluster into their own individual groups. **B,** Heatmap of differentially expressed genes (DEGs) among CardioClusters and parental cells. Selected DEGs for each group are color-coded and shown on the right. **C,** Violin plots of expression distribution for selected DEGs. **D,** Gene Ontology (GO) term analysis of molecular functions that are enriched based on DEGs. **E-F,** DEGs compared to genes expressed by freshly isolated mouse cardiac interstitial cells (mFreshCIC) represented in a Venn diagram **(E)** and heatmap of the gene set coexpressed by freshly isolated cells in comparison to CardioCluster and parental cell populations **(F).** hParental indicates human parental cells; hCardioCluster, human CardioCluster cells; BAMBI, BMP and activin membrane bound inhibitor; LIF, leukemia inhibitory factor; CXCL12, C-X-C motif chemokine 12 or stromal cell-derived factor 1 (SDF1); ACTA2, smooth muscle alpha (a)-2 actin; C. Cluster, CardioCluster.

Cellular identities of the three constituent parental lines comprising a CardioCluster are consistent with detected transcripts in each cell type. EPCs highly express ECSCR, ESM1, EGFL7 and RAC2, which are endothelial–related genes important for neovasculature and the angiogenic response (Figure 2B and 2C). The highly specific marker vascular endothelial statin (VE-statin) referred to as EGFL7, exhibits near-exclusive expression and action upon endothelial cells^55^ and is highly expressed in EPCs (Figure 2B). In contrast, many secreted angiogenic signaling molecules including vascular endothelial growth factor (VEGF) are expressed by nonendothelial cell types such as fibroblasts. MSC-enriched transcripts include the cell surface marker THY1 (also referred to as CD90) as well as Smooth muscle α-2 actin (ACTA2) (Figure 2B and 2C). Gene ontology analysis of MSCs reveals expression of extracellular matrix and adhesion molecules such as COL1A2, TIMP3 and FN1 (Figure 2B and 2D). Consistent with preference for hypoxia, MSCs are enriched for HIF1A, a transcription factor that plays a key role in response to hypoxic stimuli (Figure 2B). Lastly, transcripts associated with cell proliferation and anti-apoptotic activity such as BIRC5 and HMGB1 are differentially expressed in cCICs, as well as the chemotactic signaling molecule CXCL12, commonly referred to as stromal derived factor-1 (SDF-1), and developmental genes DKK1 and FGF5 (Figure 2B). Collectively, these data highlight that the 3 parental populations are distinctly different from one another, with EPCs and MSCs expressing endothelial and stromal-associated genes.

Transcriptome profiling of CardioClusters by scRNA-seq reveals several features distinct from the three parental cell lines. CardioClusters are enriched for transcripts in multiple categories including stem cell-relevant factors (KLF4, LIF, JAK1, SMAD7, BAMBI, NOTCH3), adhesion/extracellular-matrix molecules (integrin-α2, laminin-γ1, type 1 collagen-α1, BMP1, MMP2), and cytokines (SOD2, SDF-1, FGF2). These aforementioned DEGs were similarly enriched in freshly isolated cardiac interstitial cells (Figure 2B-D, 2F, and Supplemental Table 5), suggesting that CardioClusters adopt a transcriptome profile with features reminiscent of cardiac interstitial cells present in the myocardium rather than cultured cells. Indeed, 89 out of the 448 DEGs present in CardioClusters are also found in freshly isolated cardiac interstitial cells (Figure 2E and 2F, Supplemental Table 5), in stark contrast to the overlap with the 2D-cultured parental cell lines where only 7 DEGs are shared with freshly isolated cardiac interstitial cells (Figure 2E and 2F, Supplemental Table 5). This finding supports that standard tissue culture causes expanded cells to lose their identity, unlike cells within a 3D microenvironment. Consistent with this observation, the size of cells grown within a CardioCluster were smaller relative to 2D cultured parental counterparts (p<0.01; Supplemental Figure 6A) resembling a size more similar to freshly isolated cells, consistent with cellular remodeling that occurs alongside topological changes^56^. Thus, the 3D microenvironment of a CardioCluster promotes a more native phenotype similar to endogenous or freshly isolated cardiac cells.

### CardioClusters exert protective effects with serum starvation *in vitro* assay

Protective effects mediated by cCICs and MSCs are conferred upon serum starved neonatal rat cardiomyocytes (NRCMs) in co-culture^57^. Similarly, beneficial effects mediated by CardioClusters were assessed by co-culture with NRCMs in serum depleted conditions relative to effects conferred by cCIC, EPCs, MSCs, and a combined mixture of cCIC+EPC+MSC (C+E+M) (Figure 3A). NRCMs maintained in low serum (0.5%) were smaller relative to NRCMs maintained in high serum condition (10%) (Figure 3B and 3C, Supplemental Figure 7). CardioCluster co-culture with low serum treated NRCMs restored cardiomyocyte size within 24 hours relative to all other treatments (p<0.05; Figure 3B and 3C, Supplemental Figure 7) and also increased mRNA expression for *Desmin,* a muscle-specific type III intermediate filament protein (p<0.001; Figure 3D). Furthermore, CardioCluster co-culture increased mRNA for *Sdf-1* (p<0.05; Figure 3E), a cardioprotective cytokine and chemotactic factor for MSCs that plays an additional role in recruitment of EPCs important for angiogenesis^58, 59^. Importantly, CardioClusters offered significantly greater protection upon NRCM than actions exerted by any individual parental population (cCICs, EPCs, MSCs) or the combined C+E+M mixed population. Collectively, these results demonstrate superior protective effects of CardioClusters for NRCM in response to serum starvation challenge.

**Figure 3.**
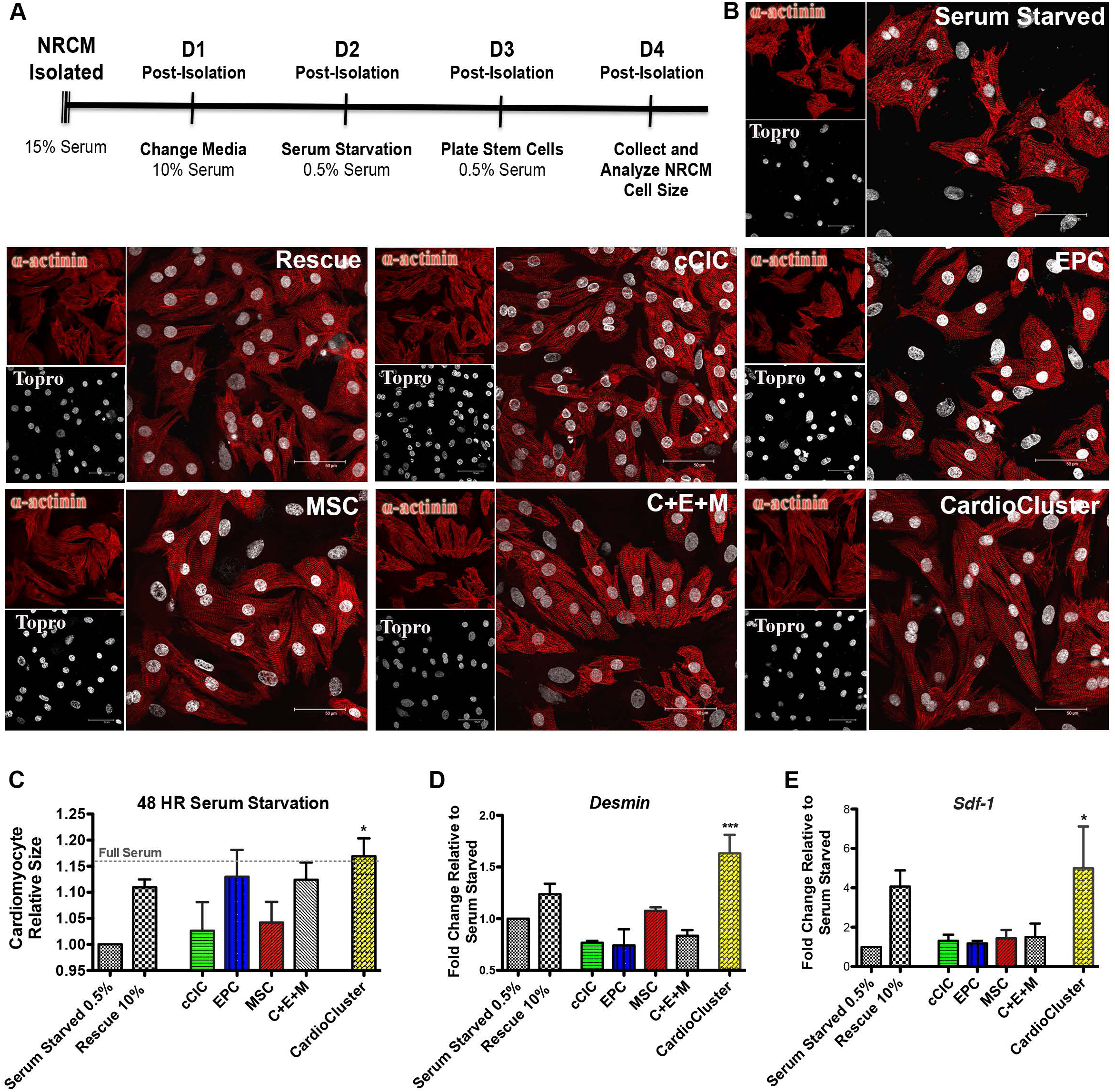
CardioClusters protect cardiomyocytes against low serum conditions *in vitro*. **A,** Timeline for neonatal rat cardiomyocyte (NRCM) low serum assay over a 4-day (D) period. **B,** Representative images of NRCM conditions: serum starved (48 hours of 0.5% serum), rescued (24 hours of 0.5% serum with 10% serum added for additional 24 hours), and experimental groups (24 hours of 0.5% serum with addition of either cCIC, EPC, MSC, C+E+M, or CardioCluster for additional 24 hours). Cardiomyocytes visualized by staining with sarcomeric actinin (α-actinin; red). TO-PRO-3 iodide (Topro; white) used to visualize nuclei. Scale bar, 50 µm. **C,** Quantitation of cardiomyocyte size relative to serum starved control. **D-E,** Gene expression of *desmin* **(D)** and *sdf-1* **(E)** in cardiomyocytes with and without the addition of cells. Data are presented as 1-way ANOVA, *p<0.05 and ***p<0.001, versus serum starved control (n=3).

### Paracrine gene expression is increased in CardioClusters after NRCM *in vitro* co-culture

Paracrine factor action is considered a primary mechanism for cardioprotective^60^, so mRNA transcript level expression for growth and immunomodulatory factors was assessed after CardioCluster co-culture for 5 days with serum depleted NRCM (Supplemental Figure 8A). mRNA levels for CardioClusters, parental cells, and the C+E+M mixed population were measured by separating fluorescently tagged cells away from the NRCM population using flow cytometric sorting (Supplemental Figure 7). mRNA transcript levels for *insulin-like growth factor* (*IGF*) and *interleukin-6* (*IL-6*) were highly elevated in CardioClusters co-cultured with NRCMS relative to any of the individual parental population (cCICs, EPCs, MSCs) or the combined C+E+M mixed population (p<0.001 and p<0.05 respectively, versus cCIC; Supplemental Figure 8B and 7C). *IGF* exerts chemotactic and growth-stimulatory effects^60^ in addition to anti-apoptotic properties^61–63^. Early release of anti-inflammatory cytokines such as *IL-6* after acute cardiac damage has been shown to be beneficial by signaling protective responses in local tissue and initiating wound healing^64^. Additionally, the cardioprotective cytokines *SDF-1* and *hepatocyte growth factor* (*HGF*), both trended towards increased expression in CardioClusters following co-culture experiments (Supplemental Figure 8D and 8E). *HGF* stimulates cell proliferation, motility, morphogenesis, angiogenesis and importantly tissue regeneration^63, 65^. Collectively these results show that at the transcript level CardioClusters induction of paracrine factors *IGF* and *IL-6* exceeds that of parental cell populations or C+E+M group when co-cultured with serum depleted NRCMs.

Several mRNAs associated with lineage specification were analyzed following co-culture of CardioClusters or parental cell populations with NRCMs. *GATA4* showed the highest expression in cCIC co-culture. Predictably, EPCs displayed the largest induction of endothelial marker *CD31*, whereas MSCs induced *SMA* gene expression after 5 days of co-culture with NRCMs (Supplemental Figure 8, F-H). Neither *CD31* nor *SMA* were significantly upregulated in CardioCluster group (Supplemental Figure 8G and 8H).

### CardioClusters are resistant to oxidative stress by *in vitro* assay

CardioClusters were substantially more resistant to cell death induced by 4 hours of H_2_O_2_ treatment after overnight low serum culture (Figure 4). Dying cells were divided into groups of early apoptosis, late apoptosis, and necrosis based on Annexin V and Sytox Blue staining (Supplemental Figure 9). CardioClusters showed significantly fewer cells in necrosis relative to any individual parental population (cCICs, EPCs, MSCs) (p<0.05 versus cCIC; Figure 4B and Supplemental Figure 9F) as visually evidenced by fewer cells rounding up and detaching from the tissue culture dish (transparent arrows; Figure 4C and 4D). These results demonstrate superiority of CardioClusters to survive oxidative stress challenge *in vitro*.

**Figure 4.**
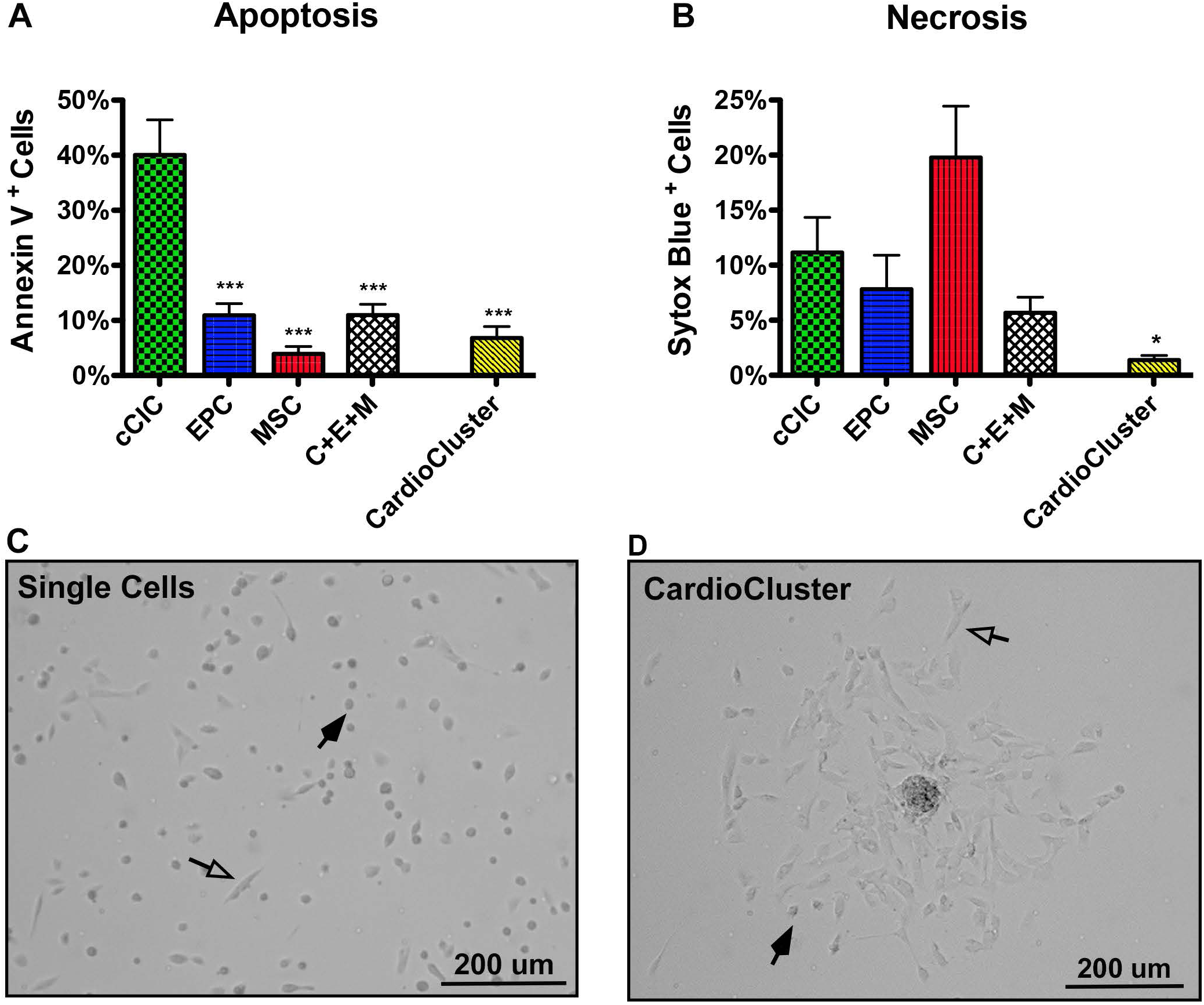
CardioClusters exhibit increased protection from oxidative stress. **A-D,** Cell death assay performed on cardiac cell populations under 24 hours of low serum (75% serum reduction), followed by 4 hours of treatment with 30 µM H_2_O_2_ in low serum medium. **A-B,** Apoptosis visualized by Annexin V **(A)** and necrosis visualized by Sytox Blue **(B).** Data are presented as 1-way ANOVA, *p<0.05, **p<0.01, ***p<0.001, versus cCIC (n=3-5). **C-D,** Brightfield images of single cells **(C)** and CardioClusters **(D)** 4 hours after H_2_O_2_ treatment. Transparent arrows highlight adherent (healthy) cells and black arrows highlight cells that have rounded up and detached from the tissue culture surface, presumably undergoing cell death.

### CardioClusters improve myocardial structure and function following infarction injury

Therapeutic efficacy of CardioClusters was assessed in a murine experimental myocardial infarction injury model of permanent coronary artery occlusion. Xenogenic human cell treatment into NOD^SCID^ recipient mice was performed at the time of infarction with direct comparison between CardioClusters and the C+E+M combined population group administered as a single cell suspension mixture. Myocardial structure and function were assessed by parasternal long axis echocardiography for four experimental groups: non-injured sham, CardioCluster, C+E+M, and vehicle-treated (Figure 5A). All groups had comparable reduction in cardiac function at 1 week post injection (wpi) demonstrating consistency of infarction injury (Figure 5, B-D, F-G and Supplemental Table 6) with average EF for all infarcted groups of approximately 30% (CardioCluster, 27±2.9%; C+E+M, 32±2.2%; Vehicle, 29±2.0%; Supplemental Figure 10). The CardioCluster-treated group showed significant cardiac functional improvement starting 4 wpi, which was sustained throughout the 20-week time course, with increased fractional shortening (FS; Figure 5B) and ejection fraction (EF; Figure 5C) versus C+E+M treatment 4 and 8 wpi and was significant at study completion for EF. In comparison, EF and FS improvements in the C+E+M treated group only began to appear at 12 and 16 wpi, respectively (Figure 5B and 5C). Terminal EF measurements at 20 wpi show EF value is highest in the CardioCluster group relative to vehicle only or C+E+M groups (40±1.9% versus 16±1.0% or 30±4.5%, respectively; Supplemental Figure 10, B-D). CardioCluster treatment shows a 45±7% improvement in EF at study completion relative to the starting value at week 1. In contrast, EF at study completion relative to week 1 for the C+E+M or vehicle only-treated groups decreased by 5±14% and 46±3%, respectively (Figure 5E). Furthermore, CardioCluster treatment group exhibits significantly smaller left ventricular internal diameter both in systole (LVID;s) and diastole (LVID;d), as well as reduction in LV end systolic and diastolic volumes (LV Vol;s and LV Vol;d). Heart rate was not significantly different among treatment groups (Figure 5, F-G and Supplemental Figure 11, A-C). Structural and functional data are detailed in Supplemental Table 6.

**Figure 5.**
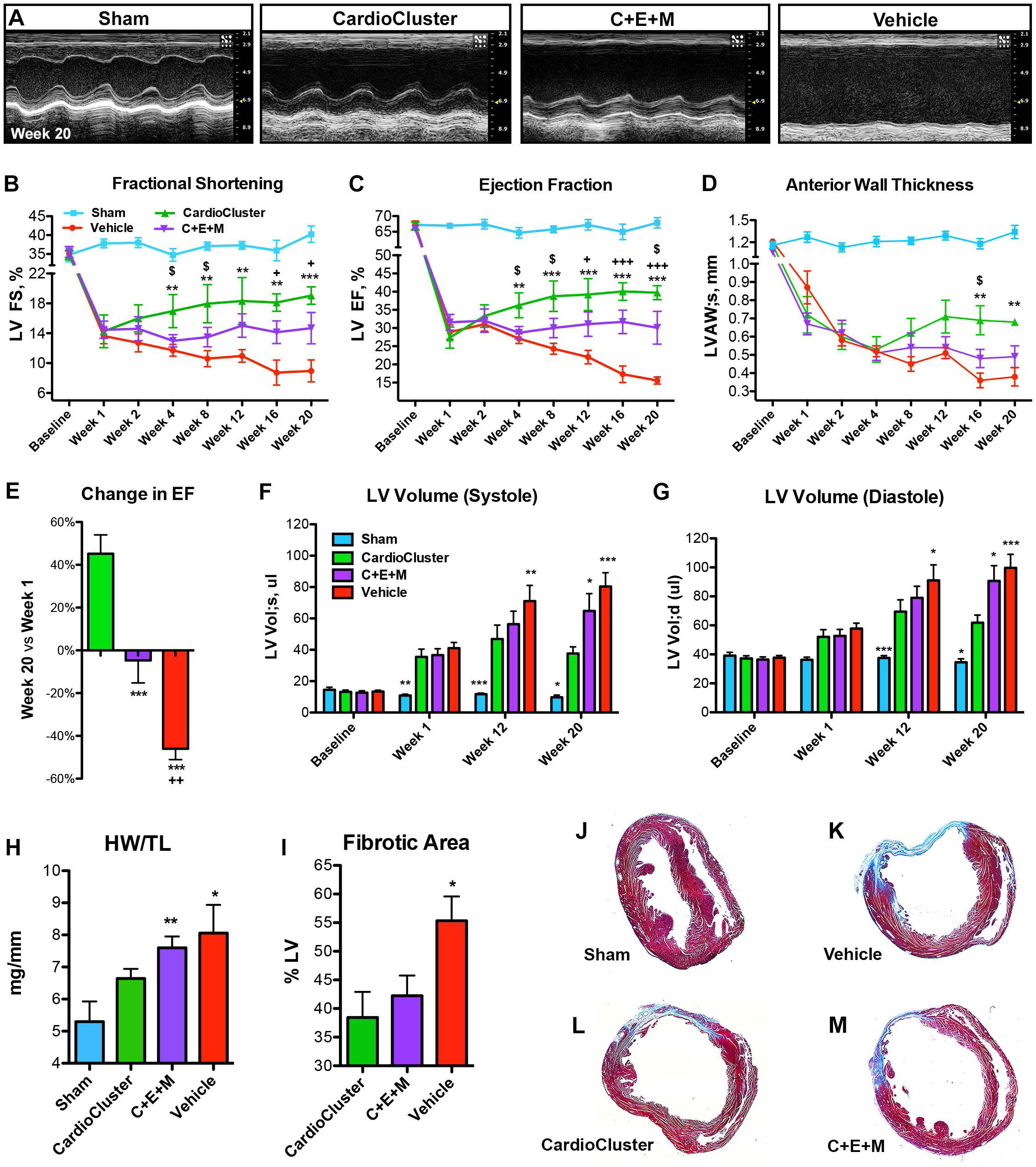
CardioCluster treatment improves left ventricular wall structure and cardiac function after myocardial injury. **A,** Representative 2D echocardiography images (M-mode) at study completion (week 20). Para-sternal short-axis view showing LV anterior wall and posterior wall movement. **B-D,** Longitudinal assessment of LV fractional shortening (FS, %) **(B),** ejection fraction (EF, %) **(C),** and anterior wall thickness in systole (LVAW;s, mm) **(D)** over 20 weeks. Data are presented as 1-way ANOVA, analyzed per timepoint, **p<0.01 and ***p<0.001, CardioCluster versus vehicle. ^+^p<0.05 and ^+++^p<0.001, C+E+M versus vehicle. ^$^p<0.05, CardioCluster versus C+E+M. Sample size specified in Supplemental Table 6. **E,** Percent change in EF from week 1 to week 20. Data are presented as 1-way ANOVA, ***p<0.001, versus CardioCluster. ^++^p<0.01, versus C+E+M (n=7-8 mice per group). **F-G,** Bar graph showing LV volume in systole (Vol;s, µl; **(F)** and in diastole (Vol;d, µl; **(G).** Data are presented as 1 Way ANOVA, *p<0.05, **p<0.01, ***p<0.001, versus CardioCluster. Sample size specified in Supplemental Table 6. **H,** Heart weight to tibia length ratio (HW/TL; mg/mm) at week 20. Data are presented as 1-way ANOVA, *p<0.05, **p<0.01, versus sham (n=4-5 mice per group). **1-M,** Masson’s Trichrome staining used to evaluate LV fibrotic area. Percentage of fibrotic LV in CardioCluster, C+E+M and vehicle-treated hearts **(I).** Data are presented as 1-way ANOVA, *p<0.05, versus CardioCluster (n=4-5 mice per group). Representative histology sections of sham **(J),** vehicle **(K),** CardioCluster **(L),** and C+E+M **(M)** treated hearts 20 weeks after Ml. Collagen-rich areas (scar tissue) are colored in blue and healthy myocardium in red.

Speckle-tracking based strain analysis is a highly sensitive echocardiographic technique for assessing left ventricular (LV) function^66, 67^. LV function was similarly reduced in all infarcted mice at 1 wpi (Figure 6A). Progressive changes consistent with adverse ventricular remodeling occur in vehicle-treated animals in agreement with conventional echocardiographic measures of function (Figure 5). LV systolic deformation in the CardioCluster-treated group showed significant improvement starting at 8 wpi and progressing through 20 wpi compared to vehicle-treated animals (p<0.001; Figure 6, B-D). Radial strain measurements at 8 and 20 wpi confirmed significant functional benefit provided by CardioCluster treatment versus 2D cultured parental C+E+M mixed population (p<0.01; Figure 6C). Peak longitudinal strain was improved in the C+E+M treated group versus vehicle alone (p<0.05; Figure 6D). Regional strain measurements assessing the area of injury further demonstrate significant improvement in LV function for CardioCluster-treated animals (Figure 6E and 6F). In the area of injury, the absolute difference in radial strain (week 1-to-week 20) for CardioCluster-treated group was 7.59±1.28% which was significantly improved relative to C+E+M and vehicle-treated groups (−1.56±1.96% and −1.88±1.61% respectively, p<0.01 versus CardioCluster group, Figure 6F). Similar improvement was seen for absolute difference in longitudinal strain 20 wpi.

**Figure 6.**
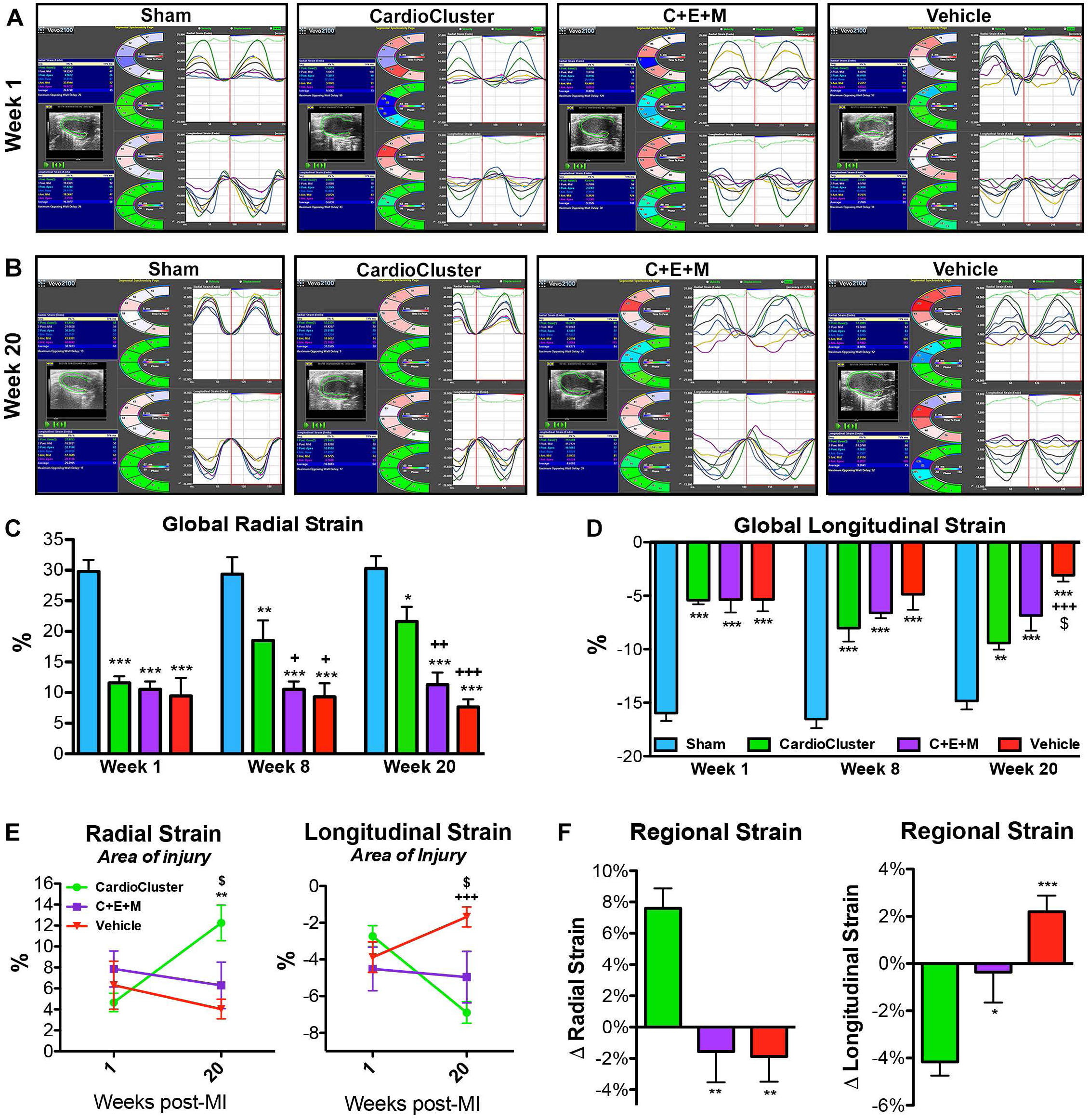
Impact of CardioClusters on cardiac function measured by cardiac strain. **A-B**, Representative images of long-axis echocardiography recording (left panel), with cross-sectional segment synchronicity map (middle panels), and radial and longitudinal strain curves (right panels) at week 1 **(A)** and week 20 **(B).** Strain curves (representing strain measures over time) are generated for the 6 standard myocardial regions, with a 7^th^ line (shown in black) denoting the average strain. **C-D,** Speckle tracking echocardiography analysis used to determine global peak radial strain (%) **(C)** and global peak longitudinal strain(%) **(D)** at week 1, week 8, and week 20. Data are presented as 2-way ANOVA, *p<0.05, **p<0.01, ***p<0.001, versus sham. ^+^p<0.01, ^++^p<0.05, ^+++^p<0.001, versus CardioCluster. ^$^p<0.05, versus C+E+M (n=6-8 mice per group). **E,** Speckle tracking echocardiography analysis used to determine regional peak radial and longitudinal strain (%) in the area of injury at week 1 and week 20. Data are presented as 2-way ANOVA, **p<0.01, versus vehicle. ^+++^p<0.001, versus CardioCluster. ^$^p<0.05, versus C+E+M (n=6-9 mice per group). **F,** Total change in regional radial and longitudinal strain from week 1 to week 20 in the area of injury. Data are presented as 1-way ANOVA, *p<0.05, **p<0.01, ***p<0.001, versus CardioCluster (n=6-9 mice per group).

CardioCluster superiority for restoring myocardial structure and function relative to the mixed population C+E+M is further reinforced by tissue morphometry and hemodynamic measurements. Cardiac hypertrophy was not a contributing factor to increasing anterior wall thickness (AWT) at 20 weeks in the CardioCluster-treated group (Figure 5D) as heart weight to tibia length ratios did not increase (HW/TL; Figure 5H) relative to the sham-operated control. In contrast, a significant increase in HW/TL is present in both vehicle control as well as C+E+M treatment groups. Fibrotic area is significantly smaller in CardioCluster-treated mice compared to vehicle at 20 wpi (38.4±4.5% of CardioCluster LV versus 55.3±4.3% of vehicle LV; Figure 5I-M) although infarct size is not significantly different between hearts receiving C+E+M or CardioClusters. Invasive hemodynamic measurement validates functional superiority of the CardioCluster-treated group showing significantly improved developed pressure over time (dP/dT) versus vehicle (Supplemental Figure 11D), in addition to increasing left ventricular developed pressure (LVDP) and P_max_-P_min_ (Supplemental Figure 11E). Collectively, these findings are evidence that CardioClusters offer significantly greater benefit for restoration of myocardial performance in this murine myocardial infarction injury model.

### CardioClusters engraft and persist in the myocardial wall following intramyocardial injection

Characteristics of CardioClusters including multicellular 3D architecture and enhanced survival are attractive features to mediate increased persistence following delivery compared to dissociated single cell suspensions such as the C+E+M mixed population. CardioCluster persistence *in vivo* was longitudinally assessed over a 20-week period by confocal microscopy (Figure 7). CardioCluster localization was tracked with co-injection of FluoSpheres tracking beads in pilot studies to confirm delivery location in tissue sections (Figure 7A). Cryosectioned hearts allowed for direct visualization of fluorophore tags without antibody labeling. All three constituent cell types were readily visualized in the myocardial wall, with MSCs at the center of the CardioCluster surrounded by a layer of cCICs and EPCs, similar to architecture observed *in vitro* (Figure 1). With CardioCluster localization confirmed coincident with the injection site, subsequent injections and imaging were performed without FluoSpheres tracking beads for long-term functional studies. CardioClusters were clearly visible within the myocardium at serial time points: 1 day, 3 days, 1-, 4-, 12-, and 20-weeks post injection (Figure 7, B-H). Antibody labeling confirmed CardioCluster persistence up to 20 weeks post injection (Figure 7E-H).

**Figure 7.**
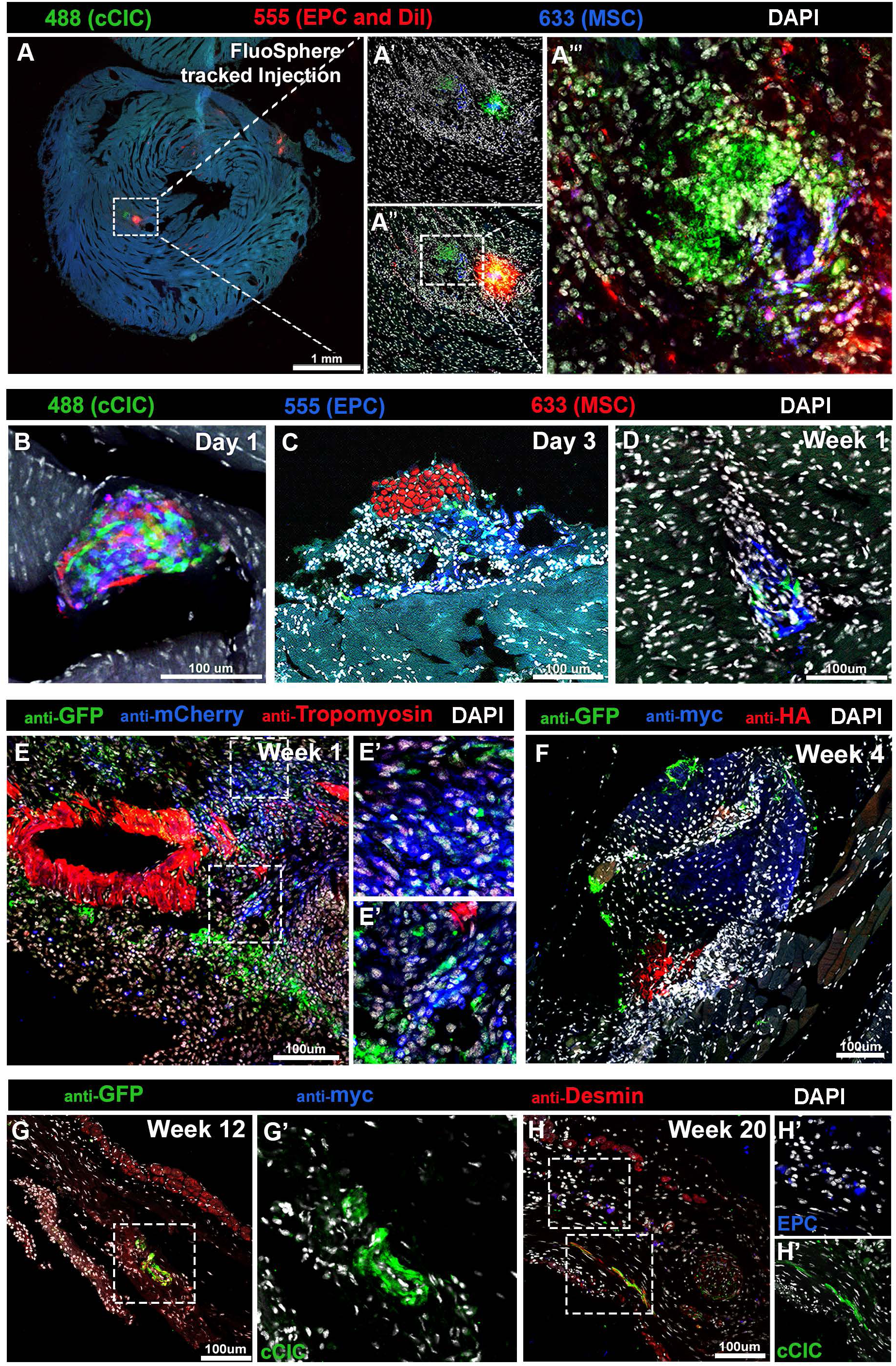
CardioCluster show enhanced engraftment and persistence in the myocardial wall. **A,** Immunofluorescent tile scan of a cryosectioned heart from an uninjured animal injected with CardioClusters day 3 post-injection tracked by FluoSpheres with no antibody labeling required for visualization of cells. **A’-A”’,** Higher magnification of areas with white dotted boxes. A’, Illustrates the removal of the 555nm channel in order to better visualize cCIC (green) and MSC (blue), without FluoSpheres. **A”-A”’,** 555nm channel restored **(A”)** and field of view magnified **(A”’). B-D,** lmmunofluorescent images from cryosectioned Ml hearts injected with CardioClusters at day 1 **(B),** day 3 **(C),** and week 1 **(D). E-H,** Antibody labeled immunofluorescent images from Ml hearts injected with CardioClusters at week 1 **(E),** week 4 **(F),** week 12 **(E),** and week 20 **(H). E’,** Higher magnification of areas with white dotted boxes. **G’,** Higher magnification of cCICs shown within white dotted box. **H’**, Higher magnification of cCICs and EPCs shown within white dotted boxes. Antibody labeling: anti-GFP labels cCIC, anti-mCherry labels EPC and MSC (E); anti-GFP labels cCIC, anti-myc labels EPC and anti-HA labels MSC (F-H). Scale bars: 1mm (A); 100µm (B-H).

### CardioClusters increase capillary density in the infarct area

Capillary density was measured in the infarct, border zone, and remote regions at 20 wpi. Non-injured controls (sham) serve as the control group compared to injured hearts (Figure 8). Notably, the CardioCluster group exhibited significantly more isolectin labeled vessels in the infarct region at 20 wpi versus both vehicle and C+E+M-treated groups (Figure 8, A and E–G). CardioCluster group capillary density in the infarct region increased 62% or 83% versus the C+E+M or vehicle only control, respectively. Within the infarct border zone at 20 wpi, both CardioCluster and C+E+M-treated groups trended toward increased capillary density versus vehicle control, but not achieving significance (Figure 8B). The remote region did not significantly increase capillary density in either CardioCluster or C+E+M groups relative to vehicle or sham (Figure 8C and 8D). Taken together, these data demonstrate a superior level of microvascularization prompted by CardioCluster treatment relative to dissociated mixed cell preparation or vehicle only control groups.

**Figure 8.**
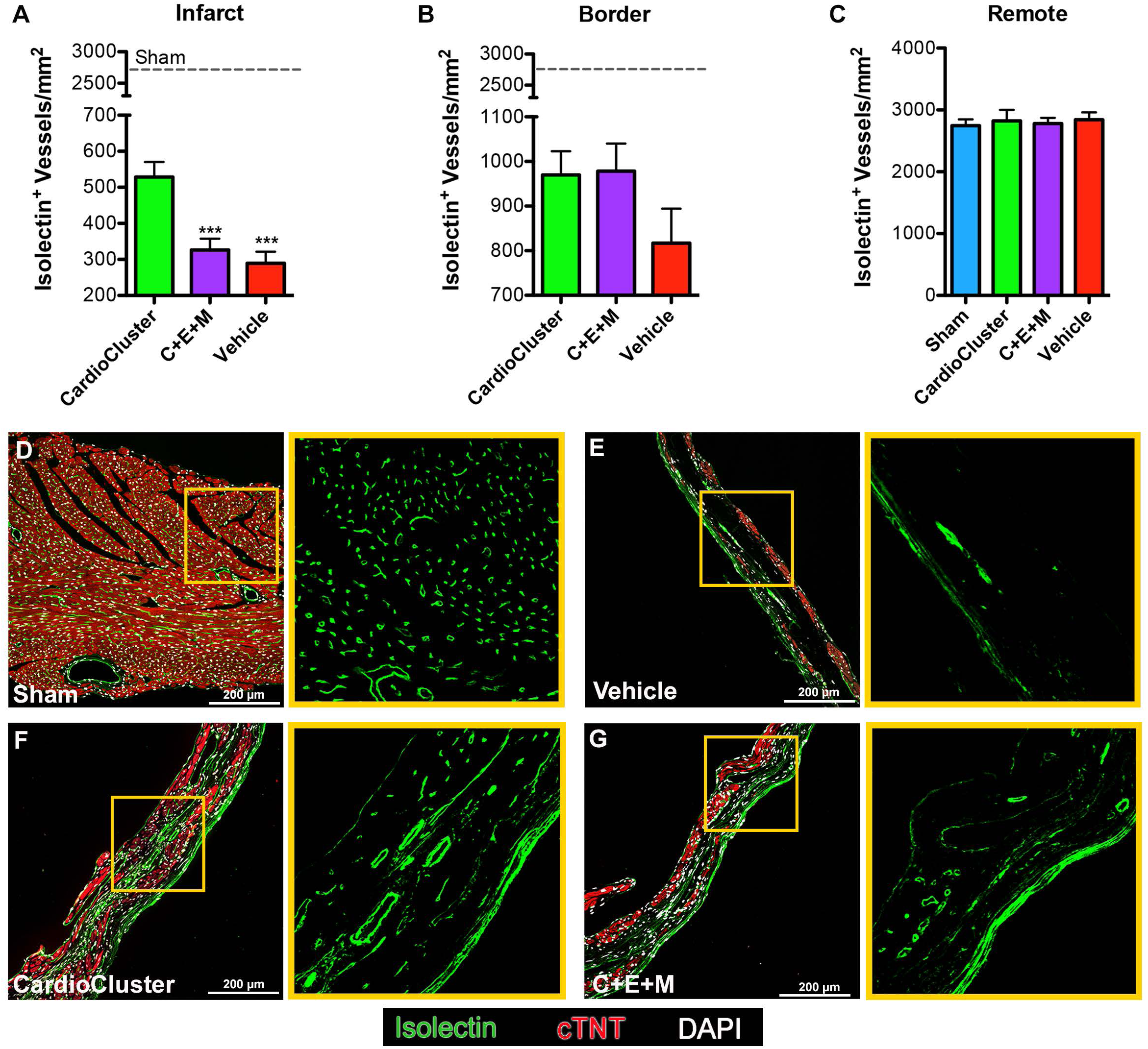
CardioCluster treatment increases capillary density in the infarct region. **A-C,** Quantitative analysis represents measurement of capillary density in the infarct **(A),** border zone **(B),** and remote **(C)** heart regions (n= 4-6 mice per group). Dashed line: mean capillary density of sham group. **D-G,** Representative image of isolectin^+^ vessels in sham **(D)** and infarct regions of Vehicle **(E),** CardioCluster **(F),** and C+E+M **(G)** used to quantitate capillaries (isolectin B4; green), cardiac troponin T (cTNT; red), and nuclei (DAPI; white). Right panels show higher magnification of isolectin^+^ vessels from areas highlighted by yellow boxes. Scale bar, 200 µm. Data are presented as 1-way ANOVA, ***p<0.001, versus CardioCluster.

### CardioCluster treatment antagonizes cardiomyocyte hypertrophy in the border and remote regions and preserves cardiomyocyte size in the infarct region

Cell therapy reduces hypertrophic remodeling following pathologic injury to blunt progression of heart failure after MI. Individual cardiomyocyte cross-sectional area was traced (Supplemental Figure 12, A-C) along with average cardiomyocyte cross-sectional area for infarct, border zone, and remote regions (Supplemental Figure 12, D-F). Treatment groups receiving either CardioClusters or the C+E+M mixed population both exhibited normalized cardiomyocyte size in the infarct region nearly identical to uninjured sham control hearts at 20 wpi (Figure 9A). Cardiomyocytes within the border zone proximal to infarction or in remote regions from the injury site were significantly smaller in the CardioCluster treatment group compared to either C+E+M or vehicle only control groups (p<0.001; Figure 9, B-C and E-G). Indeed, cardiomyocyte size in remote regions was normalized in the CardioCluster group to values similar with non-injured controls (Figure 9C and 9D). Collectively these data validate the action of CardioCluster treatment to blunt hypertrophic cellular enlargement better than dissociated mixed cell preparation or vehicle only control groups.

**Figure 9.**
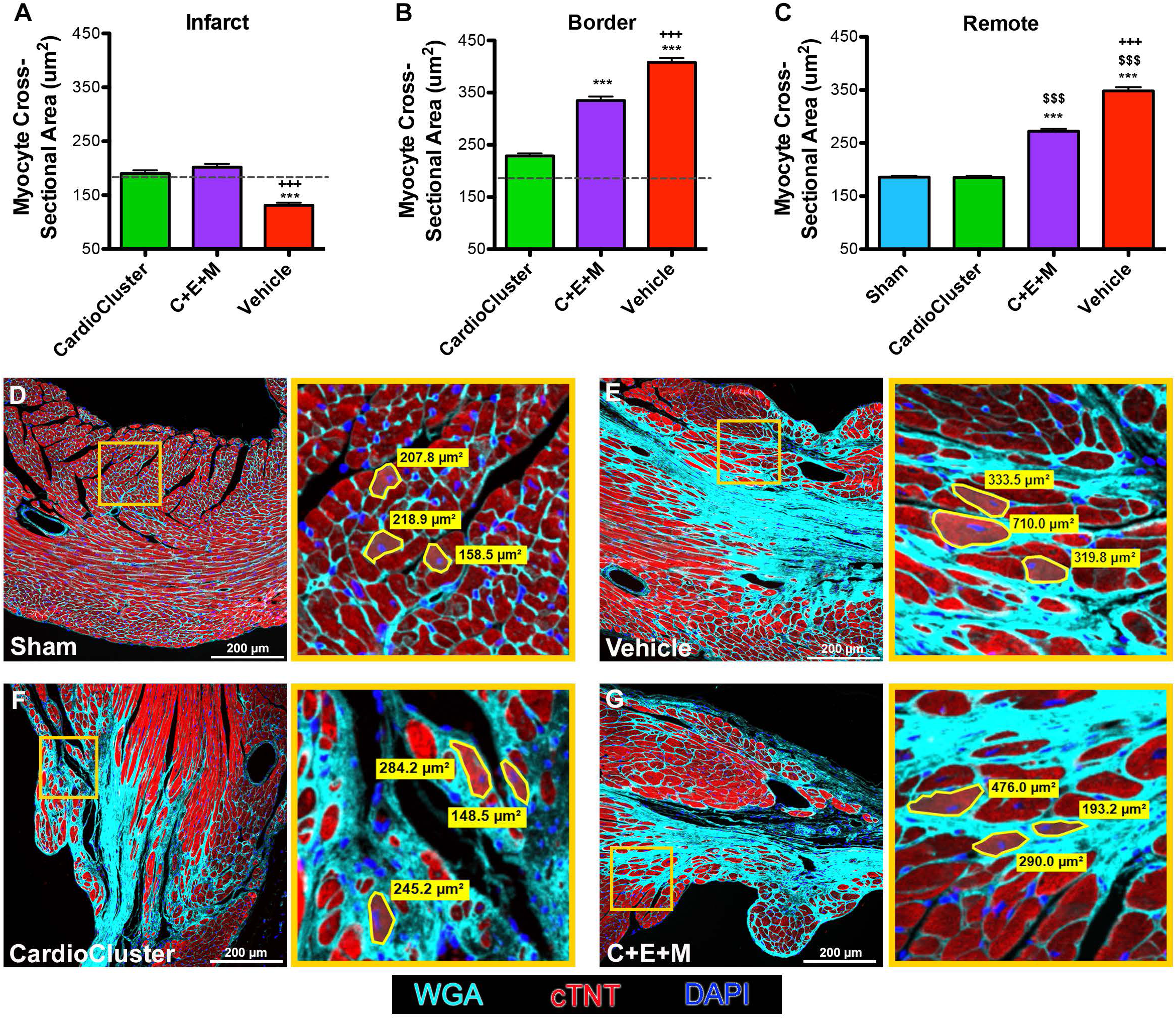
CardioCluster treatment antagonizes cardiomyocyte hypertrophy in the border and remote region and preserves cardiomyocyte size in the infarct region. **A-C,** Quantitative analysis represents measurement of cross-sectional area of cardiomyocytes from infarct **(A),** border zone **(B),** and remote **(C)** heart regions (n=4-5 mice per group). Dashed line: mean cross-sectional area of sham group. **D-G,** Representative image from sham **(D)** and border zone regions of vehicle **(E),** CardioCluster **(F),** and C+E+M **(G)** used to quantitate cardiomyocyte cross-sectional area (wheat germ agglutinin [WGA]; cyan), cardiac troponin T (cTNT; red), and nuclei (DAPI; blue). Right panels show higher magnification of areas highlighted by yellow boxes. The area of three traced cardiomyocytes per region are shown within magnified view. Scale bar, 200 µm. Data are presented as 1-way ANOVA, ***p<0.001, versus CardioCluster. ^+++^p<0.001, versus C+E+M. ^$$$^p<0.001, versus sham.

## Discussion

The preceding decade of cardiac cell therapy has produced substantial knowledge regarding optimization as well as limitations of current therapeutic interventions. In a field sometimes overshadowed by contentious debate^68–72^, it is important to remember that there are also many points of consensus. Specifically, all parties agree that inefficient cell delivery to the site of injury, low cell retention and modest efficacy of cells that do remain within the tissue are factors hampering advancement. Technical reinvention building upon prior success by incorporating ‘next generation’ approaches to surmount these established barriers represent the frontier of cell therapy research. CardioClusters introduced in this report are a novel and effective solution that integrates multiple cardiac-resident cell types into a single injectable product. The evolutionary advance offered by CardioClusters is mitigation of single cell delivery challenges through spherical self-assembly of a larger 3D structure, which provides enhanced retention to mediate repair after delivery. Inspiration for CardioClusters is drawn from prior studies showing superiority of combinatorial cell therapy^19–22^ and prolonged survival and functional engraftment of cells grown in 3D environments^73–76^ or tissue-like constructs ^27^. The multicellular structure and composition of CardioClusters represents a distinctly unique *in vitro* engineered platform to enhance the outcome of cellular therapeutics as demonstrated in preclinical testing using an established murine infarction injury model treated with a xenograft of human cardiac-derived cells.

CardioClusters were deliberately designed with multiple features anticipated to enhance efficacy. Among these properties, the essential combination of multiple cardiac cell types was enabled by our prior methodological studies to isolate and expand three distinct cardiac-derived interstitial cell types from patients with end-stage heart failure undergoing implantation of a left ventricular assist device (LVAD)^39^. End stage heart failure patients such as LVAD recipients represent likely candidates for interventional autologous therapy using cells derived from their own cardiac tissue. Combinatorial approaches harnessing beneficial attributes of multiple adult cell types are gaining acceptance as a method to enhance biological properties and efficacy based upon the tenets that: 1) no single cell population possesses all the requisite attributes for effective repair, and 2) both cardiomyogenic and non-cardiomyogenic cells contribute to myocardial repair and regeneration. Combining multiple cell types with complementary roles more efficiently mediates repair in preclinical experimental animal models of heart failure^19–22^ and is currently being assessed in the CONCERT clinical trial with patients receiving mixtures of MSCs and cardiac progenitor cells (cCIC)^77^. Similarly, induced pluripotent stem cell-derived cardiomyocytes combined with vascular cells^78^ or MSCs^79^ potentiates myocardial repair, likely due to enhanced stimulation of endogenous repair mechanisms. Efficient isolation and expansion of three distinct cardiac-resident non-myocyte populations brought together *ex vivo* to form CardioClusters is now technically feasible (Figure 1, Supplemental Figures 1 and 3) ^39^. CardioCluster biological variability depending upon the source, condition, and pathologic state of donor tissue is an important and intriguing unresolved issue to be addressed in follow-up studies based upon the proof-of-principle provided in the present report.

Another enabling feature of CardioClusters is the profound influence of aggregation upon phenotypic and biological properties of the constituent cell populations. Specifically, CardioClusters foster a transcriptional profile more consistent with freshly isolated cardiac interstitial cells compared to their monolayer counterparts (Figure 2). Even relatively short-term *in vitro* expansion of cCIC in 2D monolayer culture results in loss of identity marker gene expression and decreased population heterogeneity by single cell RNA-Seq transcriptome profiling^25^. And although cells derived from typical 2D monolayer cultures are used to seed CardioClusters, the transcriptome signature of CardioCluster cells collectively resemble each other far more than original parental cells. The CardioCluster microenvironment promotes intercellular coordination initiated within a 3D environment, unlike traditional monolayer expansion^23–25^. Increased expression of collagen type I and III, integrins (ITGA2, ITGB1, ITGA11, ITGA1, ITGAV), fibronectin, and matrix remodeling enzymes (MMP-1, MMP-2, MMP-14, TIMP-1, TIMP-2) in CardioClusters is consistent with enhanced matrix remodeling capacity of cells maintained in a 3D microenvironment^80^. Transcript data for CardioClusters also showed elevated expression of Notch3 that exerts an important regulatory role in the contexts of development and tissue regeneration for maintenance of a progenitor pool and tissue homeostasis^81^. Elevated Notch expression and superiority of 3D aggregation culture relative to conventional 2D conditions is consistent with results using pediatric cCICs cultured in 3D spheres of approximately 1500 cells, wherein 3D aggregated cCICs exert enhanced repair with increased notch signaling compared with their 2D counterparts in a right ventricular heart failure model.^50^ Restoring fresh and/or youthful characteristics to isolated cells expanded *in vitro*^82–85^ may be one way that CardioClusters provide functional benefits to the collective population.

A third enabling feature of CardioClusters is cardioprotective action, particularly under conditions of environmental stress. *In vitro* testing in co-culture assays is an established protocol to assess the potential of candidate cell types to inhibit cardiomyocyte death from pro-apoptotic challenge^57, 86^. Superior ability of CardioClusters to blunt NRCM death relative to single parental cell types supports the rationale for culturing the mixed cell population together in a 3D configuration (Figure 3). Animal studies have demonstrated that transplantation of exogenous cells exert cardioprotection through release of paracrine factors and, to a lesser extent, trans-differentiation into cardiac resident cells^79, 87–93^. This idea is supported by our transcriptomic and quantitative PCR analysis revealing the ability of cardiac interstitial cells to express a wide array of paracrine cytokines and factors that promote angiogenesis (such as VE-statin and RAC2) and cell survival (such as FGF, SDF-1, IGF, IL-6 and HGF). In vitro functional assays revealed that cardiac interstitial cells exerted prosurvival and proangiogenic effects. Furthermore, the secretome of CardioClusters also exerted protective effects on neonatal rat cardiomyocytes. Importantly, the paracrine effects of CardioClusters were more powerful than those shown with the single parental populations. These findings establish the justification for subsequent *in vivo* testing and a potential paracrine mechanistic basis for CardioCluster action.

Superior restoration of structure and function following cardiomyopathic injury is evident from comparative testing with either CardioClusters or the dissociated cell mixture of C+E+M in xenogenic treatment of NOD^SCID^ mice (Figure 5). Empirical control of CardioCluster size to <160 μm (Figure 1N and 1O) allowed for injection through a 30-gauge needle without dissociation into single cells. Improvement in FS and EF was observed starting at week 4 and maintained during the entirety of the 20-week study, with CardioCluster-treated animals showing significantly improved myocardial wall structure compared to C+E+M-treated animals concomitant with increased capillary density (Figure 8) and preserved cardiomyocyte size (Figure 9). CardioClusters persisted within the myocardial wall, with cells clearly visible up to 5 months post-injection (Figure 7). The use of human cells necessitated the use of NOD^SCID^ mice in our study. Although having a fully functioning immune system may lead to a different outcome, in view of what has been seen in previous studies the results should improve when the cells have been introduced into a model that has a working immune system. Future studies will be needed to address this point.

Meta-analysis examining cardiac stem cell (CSC) and MSC ability to treat MI in animal models found that treatment culminated in an absolute difference in EF ranging from 8-10.7% compared to control animals^94, 95^. In comparison, by 20 weeks CardioCluster treatment showed a 24.2% increase in EF compared with vehicle treatment. Our data shows the significant 2-fold improvement possible with CardioClusters versus traditional single cell therapy approaches. “Off-the-shelf” potential of CardioClusters preserved in liquid nitrogen demonstrated high viability and structural integrity indistinguishable from non-frozen counterparts (Supplemental Figure 4). Frozen/thawed CardioCluster efficacy remains to be tested *in vivo*, but the ability to mass-produce and preserve CardioClusters in frozen storage is attractive for clinical implementation planning.

The conceptual framework of CardioClusters offers almost infinite possibilities for modification and optimization for therapeutic use, as well as basic investigation of cellular interactions. For example, parameters including cell ratios, cell types, cluster size, and number of CardioClusters to inject are all worthy of further consideration. CardioCluster contain approximately 300 cells crucial for injectability through the inner diameter of a 30-gauge needle, which constrained diameter to <160 μm (Figure 1O) in murine studies. However, a larger animal could tolerate a larger gauge needle and concomitantly scaled up CardioCluster size for a greater total number of cells to be injected. With the advent of ultra-low attachment microcavity plates with between 79 – 15,000+ wells per plate, the feasibility of scaling up production is possible for future clinical application and warrants further investigation. Alternatively, ‘mini-CardioClusters’ of 50-100 cells with smaller diameter would allow a greater number of individual clusters to be injected. Ability to fine-tune CardioCluster size is a benefit distinct from traditional 3D cell aggregates such as cardiospheres where diameter is not controllable, necessitating dissociation into single cell suspensions of cardiosphere-derived cells for clinical use. Additional strategies to further enhance the CardioCluster concept could involve incorporating genetically modified cells with pro-survival factors such as Pim-1^83, 96^ or overexpression of chemokine receptor (CCR1) to enhance migration, survival and engraftment^97^. Likewise, culture condition modification using hypoxia to favor cell growth and blunt senescence-associated characteristics^98^ could dramatically alter CardioCluster biological properties. Cellular interactions occurring within 3D environments can also be tested, such as seeding the three C+E+M founder cell types together rather than sequentially, which appears to create a hollow CardioCluster (data not shown, Sussman lab). This may be attributable to EPC/MSC interaction allowing for internal cavity formation as seen during organ and tissue development^99, 100^ or may be more similar to pericyte/MSC-like endothelial cell interactions^101^. With knowledge regarding combinatorial cell therapy at a rudimentary level ^102–104^ the ‘next generation’ CardioCluster approach will benefit from further investigation given the multiple possibilities for tweaking the system to enhance the outcome.

As with any novel technological approach, there are unknowns and limitations that need to be resolved for CardioCluster development. A benefit of CardioCluster design is the quick formation time of only 48 hours from start to finish, however creation of CardioClusters necessitates that the multiple composite founder cell types must be ready for utilization in sequence within a short time frame. Since CardioClusters could be conceived using a plethora of possible cells, the time required for expansion of the parental cells may differ depending on the cell types chosen, particularly if using cells isolated from aged patients suffering from cardiomyopathic disease. Allogeneic implementation for CardioClusters might incorporate immunosuppressive agents or assembly using ‘universal’ donor cells engineered by CRISPR-mediated genome editing^105^. Prospective preparation and freeze storage of either parental cells or CardioClusters will help ease issues with timing for assembly and delivery. With respect to delivery, CardioClusters may present a safety concerns as microemboli if administered intravenously, so direct intramyocardial injection will be used, which would be the preferred approach regardless to enhance efficacy^106, 107^.

This study presents the debut of CardioClusters as a novel technical ‘next generation’ approach to improve upon established protocols using dissociated single cell preparations proven to be safe for administration to patients but of limited efficacy. Unlike other tissue engineering microfabrication approaches, the spontaneously formed CardioCluster 3D structure maximizes cellular interaction and allows for defined cell ratios, controlled size, and facilitates injectability without dissociation. This combination of features makes CardioClusters unique among current cell therapeutic approaches with demonstrated superiority over single cell mixed suspensions in mitigation of myocardial infarction damage. This initial step toward enhanced cell therapy provides a readily manipulatable platform that will benefit from further research development with the goal of potentiating cell-based therapeutic efficacy to mediate myocardial repair.

## Acknowledgments

M. Monsanto and M. Sussman designed experiments. M. Monsanto, K. White, B. Wang, Z. Ehrenberg, R. Alvarez, O. Echeagaray, S. Sengphanith, A. Muliono, N. Gude, and K. Fisher performed experiments and analyzed data. M. Monsanto and M. Sussman wrote the article. All authors read and approved the final article.

