## Supplemental Files for "Enhancing Myocardial Repair with CardioClusters"

#### SUPPLEMENTAL MATERIAL

#### Supplemental Methods

##### Human Cardiac Interstitial Cell Isolation

NIH guidelines for human research were followed as approved by IRB review (Protocol #120686). Neonatal heart tissue procured from post-mortem infants provided by a commercial source (Novogenix Laboratories) was used for isolation of human cardiac cells. Fatty tissue was excised and remaining cardiac tissue was suspended in Basic Buffer (15 mL) and minced into 1 $mm^3$ pieces. After mincing, tissue and Basic Buffer were collected in 50 mL Falcon tube. Digestive solution containing collagenase, type II 225 U/mg dry weight (Worthington, catalog #LS004174, Bio Corp, Lakewood, NJ) was dissolved in Basic Buffer (2-2.5 mg/mL) and incubated with tissue pieces for 1.5-2 hours at 37°C with continuous shaking. Digestion solution was refreshed at the one-hour time point and resulting suspensions were centrifuged at 350 *g* and resuspended in cCIC media (see Table 1). Final suspension was filtered through a 100-µm filter (Corning, Inc., catalog #352360) followed by a 40-µm filter (Corning, Inc., catalog #352340) and centrifuged at 150 *g* for 2 minutes to collect CMs. The supernatant was collected and centrifuged at 350 *g* and resuspended in cCIC media and incubated overnight at 37°C in 5% $CO_2$ incubator. The following day, cells in suspension were collected in 50 mL Falcon tube. Any cells attached were dissociated using a 1:1 mixture of Cellstripper (Corning, catalog #25-056-CI) and TrypLE Express (1X) (Thermo Fisher Scientific, catalog #12604-013). Resulting suspension was filtered through a 40-µm filter, centrifuged at 350 *g*, and resuspended in wash buffer (PBS plus 0.5% bovine serum albumin). To isolate c-Kit$^+$ cells, suspension was incubated with c-Kit–labeled beads (Miltenyi Biotec, catalog #130-091-332) and sorted according to the manufacturer's protocol. The c-Kit$^+$ fraction was divided as such: half the population was suspended in cCIC media (see Table 1) and the other half was suspended in EPC media (see Table 1). The c-Kit$^-$ population was further incubated with CD90/CD105–labeled beads and sorted according to the manufacturer's protocol (Miltenyi Biotec, catalog #130-096-253/130-051-201). Cells positive for CD90/CD105 were suspended in MSC media (see Table 1). To isolate an EPC population, at 1 week the c-Kit$^+$ population plated in EPC media was further sorted using CD133–labeled beads and sorted according to the manufacturer's protocol (Miltenyi Biotec, catalog #130-097-049). All cells were cultured at 37°C in 5% $CO_2$ incubator in their respective growth media. cCIC and EPC were split 1:2 when they reached 60-70% confluency. MSC were split 1:2 when they reached 90% confluency. All cells used in this study were mid-passage (passages 5–10).

**Flow Cytometry**

For live cell analysis, single cells were suspended in 100 µL wash buffer and incubated with primary antibody (see Table 2 for dilutions) on ice for 30 minutes. Following, cells were washed with wash buffer and incubated with secondary antibody (1:100) for 20 minutes on ice. For fixed cell analysis, cells were suspended in 4% paraformaldehyde for 5 minutes at room temperature and then washed twice with wash buffer. For c-Kit analysis requiring permeabilization, cells were washed twice and resuspended in PBS plus 0.1% Triton X-100, 0.1 M Glycine for 3 minutes, then washed once. Fixed cells were suspended in 100 µL wash buffer and incubated with primary antibody on ice for 1 hour. Following, cells were washed twice and incubated with secondary antibody (1:100) for 30 minutes on ice. For both fixed and live cells a total of 300 µl wash buffer was added post secondary incubation and the cells were analyzed by flow cytometry with a BD FACS Canto instrument. Unstained and isotype controls were used to establish baseline fluorescence levels. Data was analyzed by Flow Jo software (BD Biosciences). A minimum of 10,000 cell counts was analyzed.

**Quantitative Reverse-Transcriptase Polymerase Chain Reaction (qRT-PCR) and Bioinformatics**

Total RNA was isolated using Quick-RNA MiniPrep kit (Zymo Research, catalog #R1055) according to manufacturer's protocol. RNA concentrations were determined using a Nanodrop 2000 spectrophotometer (Thermo Fisher Scientific, catalog #ND-2000) with 500 ng concentration of RNA used to generate cDNA using an iScript cDNA Synthesis kit (Bio-Rad Laboratories, Inc, catalog #170-8891). The amplified cDNA was diluted at a ratio of 1:100 in DNase- and RNase- free water. Reactions were prepared in triplicate using 6.5 µL cDNA (equivalent to 3.25 ng total RNA) per reaction using iQ SYBER Green (Bio-Rad Laboratories, Inc, catalog #170-8882) on a CFX Real-Time PCR Detection System (Bio-Rad Laboratories, Inc, catalog #1855201). Samples were normalized to 18S and data were analyzed by $\Delta\Delta$Ct method. Primer sequences are listed in Supplemental Table 3.

**Cell and CardioCluster Morphology Measurement**

Cardiac cell populations and CardioClusters were imaged using a Leica DMIL inverted tissue culture phase contrast microscope. Morphology was measured by tracing the outline of the cells or clusters using Image J software. The three measurements analyzed were area, roundness, and length-to-width (L/W) ratios. L/W ratios were calculated by dividing feret/min feret measurements. For cell morphology, a minimum of 30 cells were measured per cell line.

**Cell Proliferation Assay**

Cell populations were plated in quadruplicate (2,000 cells/well) in a 96-well black flat bottom plate with 100 µL/well of their respective growth media. Cell proliferation rate was determined using a CyQUANT Direct Cell Proliferation Assay (Thermo Fisher Scientific, catalog #C35011) on days 0, 1, 3 and 5.

**Matrigel Tube Formation**

Growth factor reduced matrigel (Corning, catalog #356231) was used to coat a 96-well flat bottom plate (50 µl/well) and incubated for 30 minutes at 37°C. Cell populations were plated in duplicate (5,000 cells/well) suspended in 100 µL per well of EPC basal medium (see Table 1) and incubated at 37°C in $CO_2$ incubator. Images of tubular networks were acquired using a Leica DMIL inverted tissue culture phase-contrast microscope 12 to 16 hours after plating.

**Lentiviral Constructs and Cell Transduction**

All cells used for CardioCluster formation were modified by expression of fluorescent-peptide fragment tags using a 3rd generation lentiviral vector. cCICs were modified with Lenti-PGK-eGFP (Addgene), MSCs were modified with PGK-Neptune-3XHA, EPCs were modified with PGK-mOrange-3Xmyc, all at multiplicity of infection (MOI) 25. Plasmid pLenti-PGK-eGFP was used as a backbone to sub-clone pLenti-PGK-mOrange-3xmyc and pLenti-PGK-Neptune-3xHA.

**Cell Death Assay**

Human cells were plated in a 6-well dish (30,000 cells per well) and incubated in starvation media (75% FBS depleted media) with 1% PSG for 24 hours. The cells were then treated with 40μM hydrogen peroxide for 4 hours. Cells were dissociated and labeled with Annexin V (BD Biosciences; 1:175) and Propidium Iodide (PI;10mg/ml) or Sytox Blue (Life Technologies; 1:1,000) to detect apoptosis and necrosis, respectively, by flow cytometry. Data was acquired on a BD FACSAria instrument (BD Biosciences) and analyzed with FACS Diva 3 software (BD Biosciences).

**CardioCluster Preservation in Liquid Nitrogen and Viability Testing**

CardioClusters were collected, centrifuged at 150 *g* for 2 minutes, resuspended in cold freezing medium (10% DMSO in growth medium), aliquoted into cryogenic storage vials, and frozen in an isopropanol chamber stored at –80°C overnight. The following day vials were transferred to liquid nitrogen for a minimum of 24 hours prior to thaw and cell death analysis with propidium iodide (10mg/ml). Data was acquired on a BD FACSAria instrument (BD Biosciences) and analyzed with FACS Diva 3 software (BD Biosciences).

**Co-culture of Neonatal Rat Cardiomyocytes (NRCMs) with Human Cardiac Cells**

Neonatal rat hearts were excised and scissor minced prior to enzymatic digestion. Isolated NRCMs were plated in M199 media (Thermo Fisher Scientific, catalog #21157-029) with 15% FBS (Omega Scientific, Inc., catalog #FB-01) at a density of 200,000 cells per well of a 6-well culture dish. The following day, myocyte cultures were washed with PBS and incubated in M199 with 10% FBS for 24 hours. The next morning, the cells were subjected to serum starvation (0.5% FBS in M199) for 24 hours. After low serum conditions, human cardiac cells were added to the plate at a ratio of 1:10 (cCICs, EPC, MSCs, all 3 single cells combined [C+E+M], and CardioClusters) and allowed to incubate with NRCMs for an additional 24 hours in low serum conditions. Controls for NRCMs included leaving cells in 0.5% FBS M199, adding back 10% FBS M199 (Serum Rescue) or maintaining NRCMS in 10% FBS M199 for the duration of the experiment. NRCM size was visualized by staining cardiomyocytes with sarcomeric actinin (Sigma-Aldrich; 1:100 dilution) and nuclei with TO-PRO-3 iodide (Molecular Probes; 1:10,000 dilution). NRCM relative size was measured using forward scatter on a BD FACSAria instrument (BD Biosciences). Separation of NRCMs and human cardiac cells was accomplished with fluorescent cell sorting of negative cells (NRCMs) versus eGFP+, mOrange+ or Neptune+ cells. After sorting, cells were centrifuged and resuspended in RNAse buffer for isolation and quantitation of mRNA from NRCMs or human cells.

**Echocardiography and Speckle-Tracking Based Strain Measurement**

Transthoracic echocardiography was performed on lightly anesthetized mice under isoflurane (1.0-2.0%, Abbot Laboratories) using a Vevo 2100 (VisualSonics). Hearts were imaged in the 2D parasternal short-axis (SAX) view, and M-mode echocardiography of the mid-ventricle was recorded at the level of papillary muscles to calculate fractional shortening (FS). From the recorded M-mode images the following parameters were measured: left ventricular (LV) anterior wall thickness (AWT), LV posterior wall thickness (PWT), LV internal diameter (LVID), and LV volume in diastole (index: d) and systole (index: s). An LV-Trace of hearts imaged in the 2D parasternal long-axis (PLAX) view was performed in B-mode to calculate ejection fraction (EF).

Strain analysis was conducted using a speckle-tracking algorithm provided by VisualSonics (VevoStrain, VisualSonics). In brief, B-mode loops were selected from echocardiographic images based on adequate visualization of the endocardial border. A minimum of 3 consecutive cardiac cycles was selected for analysis based on image quality. Semi-automated tracing of the endocardial and epicardial borders were performed and then corrected as needed to achieve good quality tracking throughout each cine loop. Tracked images were then processed for strain measurements. Strain measurements were averaged over time resulting in curvilinear strain data points. Each long-axis view of the LV was divided into 6 standard anatomic segments for regional speckle-tracking based strain analysis throughout the cardiac cycle. Global peak strain values were averaged across all 6 segments. Regional peak strain values averaged the area of injury using segments 3, 5 and 6.

**Hemodynamic Analyses**

Invasive hemodynamic data acquisition was performed with an ADVantage PV System (ADV500, Transonic Systems Inc.) using a 1.2F PV catheter (Transonic Systems Inc., catalog #FTH-1212B-4518). Animals were sedated using 1.2 mg/mL Ketamine (VetaKet CIII, Akorn Animal Health, Inc., catalog #59399-144-10), 0.5 mg/mL Xylazine (Anased, Akorn Animal Health, Inc., catalog #59399-110-20) dosed at 10 μL/g body weight. PV catheter was pre-calibrated in 0.9% saline for at least 30 minutes at room temperature before each measurement. PV catheter was inserted through right carotid artery and advanced into LV chamber to record changes in LV pressure and volume. Hemodynamic data analysis was performed offline by LabScribe v3 software (iWorx). Mice after catheterization were immediately subjected to heart retroperfusion.

**Tissue Section Preparation**

Mice were infused with heparin (Sigma-Aldrich, catalog #H3393) at 10 U/g body weight and anesthetized using 3% chloral hydrate solution (Sigma-Aldrich, catalog #C-8383) dosed at 10 μL/g body weight. Hearts were arrested in diastole with 0.1 M CdCl2+KCl and perfused with either 1% paraformaldehydes (cryosectioned hearts) or formalin (paraffin embedded hearts) for 5 minutes at 80-100 mmHg via retrograde cannulation of abdominal aorta. Retroperfused hearts were removed from the thoracic cavity and weighed prior to fixation. Hearts were fixed overnight in either 1% paraformaldehyde at 4ºC (cryosectioned hearts) or formalin at room temperature (paraffin embedded hearts). Hearts fixed for cryosectioning were dehydrated in 30% sucrose overnight at 4°C followed by mounting in Neg50 frozen section medium (Thermo Fisher Scientific, catalog #6502) on dry ice. Tissues were cryosectioned at 20 μm thickness at -20ºC.

Formalin fixed hearts were processed for paraffin embedding and sectioned at 7 μm thickness at room temperature.

**Capillary Density Measurement**

Paraffin sections were immunolabeled with isolectin GS-IB4 conjugated to Alexa Fluor 568 (Thermo Fisher Scientific, catalog # I21412; 1:100 dilution) to visualize vasculature, in combination with cardiac troponin T conjugated to Alexa Fluor 488 (Biocompare, catalog #bs-10648R-A488; 1:200 dilution) and 4′,6-diamidino-2-phenylindole (DAPI). Scans consisted of infarct, border and remote regions for each heart analyzed. The analysis software on a Leica TCS SP8 Confocal Microscope and Image J software were used to quantitate the number of positive cells in each field of view. A minimum of 3 independent fields of view per cardiac region was measured. The area of cardiac tissue in each field of view was measured and used to normalize capillary numbers per $mm^2$. N=4-6 hearts per group measured at week 20.

**Cardiomyocyte Cross-sectional Area Measurement**

Paraffin sections were immunolabeled with cardiac troponin T conjugated to Alexa Fluor 488 to visualize cardiomyocytes, wheat germ agglutinin conjugated to 680 (Thermo Fisher Scientific, catalog #W32465; 1:500 dilution) to outline cellular membranes, and DAPI to visualize nuclei. Cardiomyocytes were measured in the infarct, border, and remote regions. Cross-sectioned cardiomyocytes with a centrally located nucleus were considered. Approximately 50 cardiomyocytes from 3 independent fields of view per heart region were measured using the SP8 TCS Leica drawing tool to trace cardiomyocyte cross-sectional area. N=4-5 hearts per group measured at week 20.

**Infarct Size Quantitation**

Trichrome Stain (Masson) Kit (Sigma-Aldrich, catalog #HT15) was used to stain for collagen deposition in sham and infarcted hearts according to manufacturer's protocol. Staining was visualized using a Leica DMIL6000 microscope using XY stage tile scan and automatically stitched by Leica LAS X analysis software. Area of live versus dead myocardium was measured using the drawing tool in the SP8 TCS Leica Software using scar length over total LV length. Tissue sections from apex to mid-wall were averaged for scar quantification. N=4-5 hearts per group measured at week 20.

#### Supplemental Figure Legends

**Supplemental Figure 1. In vitro lineage comparison to established cell lines**

**A**, Representative images of tubular network formation when plated on growth factor reduced matrigel for HUVEC, cCIC, EPC**,** and MSC. **B–E**, Bar graphs using established cell lines, HUVEC and bone marrow-derived MSC (BM MSC), to assess the potential of cardiac interstitial cells to commit to an angiogenic **(B, C)**, smooth muscle **(D)**, and cardiogenic **(E)** fate (n=FH-09 heart [cell line used to create CardioClusters], run in triplicate). Data are presented as 1 Way ANOVA, *p<0.05, **p<0.01, ***p<0.001, versus cCIC, $^{*}$excludes HUVEC and BM MSC from statistical analysis, $^{+}$includes HUVEC and BM MSC in statistical analysis. Scale bar, 200 μm. GATA4 indicates GATA binding protein 4; PECAM-1, platelet endothelial cell adhesion molecule; SMA, α-smooth muscle actin; and VWF, von Willebrand factor.

**Supplemental Figure 2. CardioCluster formation and characterization**

**A**, Human phosphorglycerate kinase (hPGK) lentiviral backbones of fluorescent protein tags used to transduce parental cell lines. cCIC express eGFP, EPC express mOrange (with a 3x myc tag), and MSC express Neptune (with a 3x HA tag). **B**, Representative flow cytometry plots showing the percentage of cells expressing their respective fluorescent proteins. **C**, CardioClusters are formed using 96 well, ultra-low attachment round bottom plates in a two-step process. The first step generates the inner core composed of cCIC and MSC, and the second step forms the outer EPC layer. The inner core of cCIC and MSC is seeded for 24 hours. The EPC are added the following day and resulting cell mixture is incubated for an additional 48-72 hours prior to experimentation. Schematic adapted from previous publication[1]. **D**, CardioCluster morphometric parameters measuring area (a.u.: arbitrary units.), roundness, and length-to-width (L/W) ratio over a 7 day time course for CardioClusters ranging from 100-1000 cells. **E**, Still frame images from a video showing CardioCluster formation.

**Supplemental Figure 3. Cell morphology measurements**

**A**, Representative phase contrast images of individually traced cells from the three cell populations isolated from human heart samples. **B-D**, Scatter plots showing individual heart averages for the morphometric parameters of length-to-width (L/W) ratio **(B)**, roundness **(C)**, and area (arbitrary units [a.u.]) **(D)** (n=3 heart samples, minimum of 30 cells traced per cell type, per patient).

**Supplemental Figure 4. CardioClusters frozen in liquid nitrogen maintain structural integrity and viability**

**A**, Representative flow cytometry plots showing propidium iodide (PI) gating strategy used in freezing assay. **B**, Brightfield images showing cell outgrowth of non-frozen versus liquid nitrogen frozen CardioClusters. **C**, Quantification of percent necrotic ($PI^+$) cells from non-frozen versus liquid nitrogen frozen experimental groups. Data are presented as 1-way ANOVA, *$p<0.05$, **$P<0.01$, ***$p<0.001$, versus non-frozen CardioClusters. ns indicates not statically significant.

**Supplemental Figure 5. Quality control for single-cell RNA sequencing**

**A**, Cell size quantification for average diameter of 2D cultured cCICs, EPCs, and MSCs versus cells cultured within a 3D CardioCluster. Data are presented as 1-way ANOVA, *$p<0.05$, **$P<0.01$, ***$p<0.001$ (n=3-4 repeated experiments). **B**, Violin plots for the number of genes, unique molecular identifiers (UMIs) and percent mitochondrial genes used for single-cell RNA sequencing quality control. **C**-**F**, Cell Ranger 2.0 quality control summary for cCIC **(C)**, EPC **(D)**, MSC **(E)** and CardioCluster **(F)**.

**Supplemental Figure 6. CardioClusters restore NRCM morphology following serum starvation**

Representative flow cytometry plots showing forward scatter (FSC-A) used to quantitate neonatal rat cardiomyocyte (NRCM) mean area. Human cardiac cell populations are excluded from analysis by gating out fluorescently tagged cells (represented by pink cells in plots). NRCMs included in analysis are represented in blue.

**Supplemental Figure 7. CardioClusters have increased paracrine and commitment gene expression after *in vitro* co-culture with cardiomyocytes**

**A**, Timeline for NRCM co-culture commitment assay. **B-H**, Gene expression in interstitial cells after a 7-day co-culture with NRCMs. **B**, *IGF* **C**, *IL-6* **D**, *SDF-1* **E**, *HGF* **F**, *GATA4* **G**, *CD31* and **H**, *SMA* gene expression (n=3 NRCM preps). Data are presented as 1-way ANOVA, *$p<0.05$, **$p<0.01$, ***$p<0.001$, versus cCIC.

**Supplemental Figure 8. Representative staining for markers of apoptosis and necrosis**

**A**, Representative flow cytometry plots showing Annexin V/Sytox Blue gating strategy used in cell death assay. **B-F**, Representative flow cytometry plots showing Annexin V/Sytox Blue labeling following cell death assay on cardiac cell populations under 24 hours of low serum (75% serum reduction) and 4 hours of treatment with 30 μM $H_2O_2$ in low serum medium for cCIC **(B)**, EPC **(C)**, MSC **(D)**, C+E+M **(E)**, and CardioCluster **(F)**.

**Supplemental Figure 9. Ejection fraction for individual mice grouped by surgery**

**A-D**, Longitudinal assessment of ejection fraction (EF, %) over 20 weeks for individual mice by surgery type: sham **(A)**, vehicle **(B)**, C+E+M **(C)**, and CardioCluster **(D)**.

**Supplemental Figure 10. Hemodynamic data confirms CardioCluster treatment preserves cardiac function**

**A-B**, LV internal diameter (LVID) in systole (LVID;s; **A**) and diastole (LVID;d; **B**). Sample size specified in Supplemental Table 6. **C,** Heart rates (beats per minute [bpm]) for sham, CardioCluster, C+E+M, and vehicle treatment groups shown at baseline, week 1, week 12 and week 20. Sample size specified in Supplemental Table 6. **D-E**, Hemodynamic analysis showing developed pressure over time (dP/dt, mmHg/sec; **D**) and left ventricular developed pressure (LVDP, mmHg) and pressure max minus pressure min ($P_{max}$-$P_{min}$, mmHg) shown at week 20 (**E**; n=3-5 mice per group). Data are presented as 1-way ANOVA, *p<0.05, versus sham.

**Supplemental Figure 11. CardioCluster treatment antagonizes cardiomyocyte hypertrophy in the border and remote region and preserves cardiomyocyte size in**

**the infarct region**

**A-C**, Scatter plots showing cardiomyocyte cross-sectional area for each individually traced cell in the infarct **(A)**, border **(B)**, and remote **(C)** heart regions. **D-F**, Individual mean for each mouse used to quantify cardiomyocyte cross-sectional area in the infarct **(D)**, border **(E)**, and remote **(F)** heart regions shown by scatter plots (n=4-5 mice per group).

**Supplemental Table 1. List of Media**

| | Component | Catalog Number |
|---|---|---|
| Cardiac Interstitial Cell Medium | F12 HAM's (1x) | SH30026.01, HyClone |
| | 10% ES FBS | 16141079, Gibco |
| | 1% Penicillin-Streptomycin-Glutamine (100X) | 10378016, Gibco |
| | 5 mU/mL human erythropoietin | E5627, Sigma-Aldrich |
| | 10 ng/mL human recombinant basic FGF | HRP-0011, Biopioneer |
| | 0.2 mM L-Glutathione | 66013-256, Sigma-Aldrich |
| Endothelial Progenitor Cell Medium | EBM-2 Basal Medium | CC-3156, Lonza |
| | EGM-2 Kit Supplements and Growth Factors:<br>• 0.5 mL Human Epidermal Growth Factor<br>• 0.5 mL Insulin-Like Growth Factor-1<br>• 0.5 mL Vascular Endothelial Growth Factor<br>• 0.5 mL HEPARIN<br>• 0.5 mL Gentamicin Sulfate Amphotericin-B<br>• 0.5 mL Ascorbic Acid<br>• 2.0 mL Human Fibroblast Growth Factor-B<br>• 2.0 Hydrocortisone<br>• 10 mL FBS | CC-4176, Lonza |
| Mesenchymal Stem Cell Medium | 10.1 g/L Minimum Essential Medium Eagle, Alpha Modification | M0644, Sigma-Aldrich |
| | 20% FBS | FB-01, Omega Scientific, inc. |
| | 1% Penicillin-Streptomycin-Glutamine (100X) | 10378-016, Gibco |
| | Cell Culture Grade Water | |
| Basic Buffer | 11 g/L Minimum Essential Medium Eagle, Joklik Modification | M0518, Sigma-Aldrich |
| | 3 mM HEPES | H3375, Sigma-Aldrich |
| | 1% Penicillin-Streptomycin-Glutamine (100X) | 10378-016, Gibco |
| | 10 mM Taurine | T0625, Sigma-Aldrich |
| | Insulin, solvate in 3% Acetic Acid/PBS | I-5500, Sigma-Aldrich |
| | 1% Amphotericin B | 15290-018, Invitrogen |
| | 50 mg Gentamicin | G1397, Sigma-Aldrich |
| | Cell Culture Grade Water | |

**Supplemental Table 2. List of Antibodies**

| Antibody | Vendor | Catalog Number | Dilution Flow | Dilution ICC/IHC |
|---|---|---|---|---|
| **C-Kit** (CD117) | R&D systems | AF1356 | 1:33 | - |
| **Thy-1** (CD90) | Biolegend | 328109 | 1:33 | - |
| **Endoglin** (CD105) | Biolegend | 323203 | 1:33 | - |
| **Prominin-1** (CD133) | Thermo Fisher Scientific | PA5-38014 | 1:33 | - |
| **PTPRC** (CD45) | Biolegend | 368507 | 1:33 | - |
| **cTNT** | Biocompare | bs-10648R-A488 | - | 1:200 |
| **Tropomyosin** | Sigma-Aldrich | T 9283 | - | 1:200 |
| **eGFP** | Molecular Probes | A-11122 | - | 1:100 |
| **mCherry** | Thermo Fisher Scientific | M11240 | - | 1:100 |
| **Isolectin GS-IB4** | Thermo Fisher Scientific | I21412 | - | 1:100 |
| **WGA** | Thermo Fisher Scientific | W32465 | - | 1:500 |
| **Myc tag** | Thermo Fisher Scientific | PA3-981 | | 1:100 |
| **HA-prope** | Santa Cruz Biotechnology | SC-7392 | - | 1:100 |
| **DAPI (**4′,6-diamidino-2-phenylindole**)** | Sigma-Aldrich | D9542 | - | 1:10,000 |
| **Phalloidin** | Thermo Fisher Scientific | A12379 | - | 1:1,000 |

**Supplemental Table 3. List of Primers**

| Target | Fwd Primer Sequence | Rev Primer Sequence |
|---|---|---|
| PECAM1 | CCAAGCCCGAACTGGAATCT | CACTGTCCGACTTTGAGGCT |
| GATA4 | CTCAGAAGGCAGAGAGTGTGTCAA | CACAGATAGTGACCCGTCCCAT |
| HGF | GGCTGGGGCTACACTGGATTG | CCACCATAATCCCCCTCACAT |
| IGF | GACCGCGGCTTCTACTTCAG | AAGAACTTGCCCACGGGGTAT |
| SMA | CCCAGCCAAGCACTGTCAGGAATCCT | TCACACACCAAGGCAGTGCTGTCC |
| CXCL12 | CAGTCAACCTGGGCAAAGCC | AGCTTTGGTCCTGAGAGTCC |
| IL-6 | TCGAGCCCACCGGGAACGAA | GCAGGGAAGGCAGCAGGCAA |
| 18S | CGAGCCGCCTGGATACC | CATGGCCTCAGTTCCGAAAA |

**Supplemental Table 4: Online link to dataset containing DEGs**

https://docs.google.com/document/d/1WEP7UBdSGIYcyRHwLfrpIZ_F_Wu9NSk9tmmm5ATLjwE/edit?usp=sharing

**Supplemental Table 5: Online link to dataset containing DEGs enriched in freshly isolated cardiac interstitial cells**

https://docs.google.com/document/d/13v1Nk2Ytv0GKM9QXDmeUSmW80Tr4E9flhhVf5GP16lw/edit?usp=sharing

#### Supplemental Table 6. Heart Rate and Echocardiographic Data

HEART RATE (bpm)

| Heart Rate | SHAM | | | CardioCluster | | | C+E+M | | | Vehicle | | |
|---|---|---|---|---|---|---|---|---|---|---|---|---|
| Week | Mean | SEM | N | Mean | SEM | N | Mean | SEM | N | Mean | SEM | N |
| Baseline | 487.95 | 15.13 | 12 | 472.26 | 15.77 | 17 | 493.17 | 14.72 | 15 | 471.78 | 17.49 | 16 |
| 1 | 449.23 | 17.07 | 12 | 470.50 | 10.91 | 12 | 471.52 | 19.70 | 15 | 426.94 | 16.35 | 16 |
| 2 | 437.72 | 28.47 | 12 | 462.27 | 18.23 | 12 | 469.08 | 13.96 | 15 | 433.02 | 22.80 | 14 |
| 4 | 439.43 | 14.62 | 12 | 488.52 | 26.94 | 12 | 503.79 | 24.43 | 15 | 445.81 | 15.40 | 16 |
| 8 | 462.07 | 16.13 | 12 | 470.52 | 14.81 | 11 | 476.45 | 20.96 | 13 | 423.57 | 19.74 | 16 |
| 12 | 492.90 | 13.07 | 11 | 469.22 | 18.92 | 11 | 507.16 | 13.92 | 13 | 507.34 | 13.78 | 11 |
| 16 | 492.68 | 15.73 | 8 | 443.20 | 20.18 | 8 | 478.72 | 26.11 | 10 | 491.36 | 22.22 | 8 |
| 20 | 456.47 | 25.34 | 6 | 438.72 | 19.99 | 7 | 445.56 | 28.76 | 7 | 504.69 | 13.35 | 8 |

ANTERIOR WALL THICKNESS; SYSTOLE (mm)

| LVAW;s | SHAM | | CardioCluster | | C+E+M | | Vehicle | |
|---|---|---|---|---|---|---|---|---|
| Week | Mean | SEM | Mean | SEM | Mean | SEM | Mean | SEM |
| Baseline | 1.16 | 0.05 | 1.16 | 0.04 | 1.07 | 0.04 | 1.21 | 0.04 |
| 1 | 1.27 | 0.07 | 0.72 | 0.10 | 0.67 | 0.06 | 0.87 | 0.09 |
| 2 | 1.13 | 0.06 | 0.60 | 0.07 | 0.62 | 0.07 | 0.58 | 0.03 |
| 4 | 1.21 | 0.07 | 0.53 | 0.07 | 0.51 | 0.04 | 0.52 | 0.03 |
| 8 | 1.22 | 0.05 | 0.62 | 0.08 | 0.54 | 0.08 | 0.45 | 0.04 |
| 12 | 1.29 | 0.06 | 0.71 | 0.09 | 0.54 | 0.06 | 0.51 | 0.03 |
| 16 | 1.18 | 0.07 | 0.69 | 0.08 | 0.48 | 0.05 | 0.36 | 0.04 |
| 20 | 1.34 | 0.09 | 0.68 | 0.02 | 0.49 | 0.06 | 0.38 | 0.05 |

ANTERIOR WALL THICKNESS; DIASTOLE (mm)

| LVAW;d | SHAM | | CardioCluster | | C+E+M | | Vehicle | |
|---|---|---|---|---|---|---|---|---|
| Week | Mean | SEM | Mean | SEM | Mean | SEM | Mean | SEM |
| Baseline | 0.75 | 0.07 | 0.76 | 0.06 | 0.66 | 0.04 | 0.81 | 0.04 |
| 1 | 0.90 | 0.06 | 0.67 | 0.10 | 0.60 | 0.03 | 0.75 | 0.06 |
| 2 | 0.83 | 0.05 | 0.53 | 0.05 | 0.53 | 0.05 | 0.54 | 0.03 |
| 4 | 0.84 | 0.05 | 0.49 | 0.05 | 0.43 | 0.04 | 0.50 | 0.03 |
| 8 | 0.79 | 0.03 | 0.49 | 0.06 | 0.46 | 0.07 | 0.44 | 0.04 |
| 12 | 0.88 | 0.04 | 0.59 | 0.06 | 0.48 | 0.05 | 0.47 | 0.03 |
| 16 | 0.84 | 0.05 | 0.56 | 0.06 | 0.38 | 0.05 | 0.28 | 0.02 |
| 20 | 0.97 | 0.06 | 0.52 | 0.03 | 0.40 | 0.06 | 0.35 | 0.03 |

POSTERIOR WALL THICKNESS; SYSTOLE (mm)

| LVPW;s | SHAM | | CardioCluster | | C+E+M | | Vehicle | |
|---|---|---|---|---|---|---|---|---|
| Week | Mean | SEM | Mean | SEM | Mean | SEM | Mean | SEM |
| Baseline | 1.23 | 0.06 | 1.39 | 0.04 | 1.50 | 0.05 | 1.32 | 0.05 |
| 1 | 1.27 | 0.08 | 1.25 | 0.07 | 1.28 | 0.12 | 1.25 | 0.09 |
| 2 | 1.29 | 0.06 | 1.22 | 0.09 | 1.18 | 0.10 | 1.26 | 0.10 |
| 4 | 1.05 | 0.09 | 1.11 | 0.06 | 1.01 | 0.09 | 1.12 | 0.07 |
| 8 | 1.08 | 0.06 | 0.83 | 0.12 | 1.00 | 0.08 | 0.96 | 0.08 |
| 12 | 1.10 | 0.06 | 1.17 | 0.09 | 1.32 | 0.09 | 0.96 | 0.08 |
| 16 | 1.16 | 0.09 | 1.13 | 0.09 | 1.06 | 0.15 | 0.99 | 0.16 |
| 20 | 1.20 | 0.07 | 1.34 | 0.10 | 1.03 | 0.06 | 1.20 | 0.15 |

POSTERIOR WALL THICKNESS; DIASTOLE (mm)

| LVPW;d | SHAM | | CardioCluster | | C+E+M | | Vehicle | |
|---|---|---|---|---|---|---|---|---|
| Week | Mean | SEM | Mean | SEM | Mean | SEM | Mean | SEM |
| Baseline | 0.66 | 0.06 | 0.82 | 0.06 | 0.78 | 0.04 | 0.86 | 0.06 |
| 1 | 0.78 | 0.04 | 0.86 | 0.06 | 0.83 | 0.09 | 0.92 | 0.09 |
| 2 | 0.83 | 0.06 | 0.87 | 0.09 | 0.77 | 0.07 | 1.00 | 0.07 |
| 4 | 0.72 | 0.08 | 0.87 | 0.05 | 0.78 | 0.07 | 0.89 | 0.06 |
| 8 | 0.75 | 0.06 | 0.62 | 0.06 | 0.85 | 0.07 | 0.79 | 0.07 |
| 12 | 0.76 | 0.05 | 0.96 | 0.07 | 1.08 | 0.07 | 0.83 | 0.06 |
| 16 | 0.81 | 0.06 | 0.88 | 0.06 | 0.86 | 0.13 | 0.83 | 0.13 |
| 20 | 0.82 | 0.09 | 1.16 | 0.10 | 0.80 | 0.04 | 1.00 | 0.14 |

LEFT VENTRICULAR VOLUME; SYSTOLE (μL)

| LV Vol;s | SHAM | | CardioCluster | | C+E+M | | Vehicle | |
|---|---|---|---|---|---|---|---|---|
| Week | Mean | SEM | Mean | SEM | Mean | SEM | Mean | SEM |
| Baseline | 14.45 | 1.66 | 13.30 | 1.18 | 12.93 | 1.38 | 13.33 | 0.90 |
| 1 | 10.81 | 0.90 | 37.98 | 5.44 | 37.63 | 4.19 | 41.07 | 3.59 |
| 2 | 12.46 | 1.23 | 42.28 | 3.22 | 45.18 | 4.15 | 45.85 | 3.16 |
| 4 | 13.22 | 1.41 | 41.83 | 5.20 | 51.40 | 4.58 | 56.60 | 6.31 |
| 8 | 11.84 | 0.59 | 45.20 | 5.41 | 54.85 | 6.06 | 64.57 | 7.43 |
| 12 | 11.71 | 0.67 | 46.84 | 8.88 | 56.34 | 8.24 | 71.03 | 10.05 |
| 16 | 11.94 | 1.64 | 40.78 | 5.58 | 56.15 | 10.42 | 71.52 | 9.07 |
| 20 | 9.72 | 1.37 | 37.64 | 4.29 | 64.76 | 11.04 | 80.37 | 8.77 |

LEFT VENTRICULAR VOLUME; DIASTOLE (μL)

| LV Vol;d | SHAM | | CardioCluster | | C+E+M | | Vehicle | |
|---|---|---|---|---|---|---|---|---|
| Week | Mean | SEM | Mean | SEM | Mean | SEM | Mean | SEM |
| Baseline | 39.15 | 2.25 | 38.20 | 1.82 | 37.01 | 2.13 | 37.72 | 1.51 |
| 1 | 36.21 | 1.82 | 52.62 | 5.80 | 53.51 | 4.75 | 57.78 | 3.73 |
| 2 | 37.96 | 1.68 | 63.20 | 3.65 | 64.71 | 4.42 | 63.42 | 3.77 |
| 4 | 36.19 | 1.95 | 63.29 | 5.73 | 70.53 | 5.35 | 75.31 | 7.11 |
| 8 | 37.20 | 1.62 | 69.70 | 5.33 | 75.39 | 6.63 | 82.94 | 7.85 |
| 12 | 37.48 | 1.65 | 69.48 | 8.12 | 78.99 | 8.06 | 91.04 | 10.67 |
| 16 | 34.83 | 2.15 | 64.71 | 7.01 | 77.37 | 10.64 | 87.73 | 9.46 |
| 20 | 34.53 | 2.42 | 61.78 | 5.26 | 90.63 | 10.58 | 99.61 | 9.46 |

EJECTION FRACTION (%)

| EF | SHAM | | CardioCluster | | C+E+M | | Vehicle | |
|---|---|---|---|---|---|---|---|---|
| Week | Mean | SEM | Mean | SEM | Mean | SEM | Mean | SEM |
| Baseline | 67.05 | 1.64 | 68.57 | 0.71 | 65.90 | 0.99 | 67.10 | 1.32 |
| 1 | 66.48 | 1.24 | 27.37 | 2.92 | 31.58 | 2.17 | 28.83 | 1.98 |
| 2 | 65.60 | 1.58 | 33.29 | 3.10 | 32.03 | 3.20 | 31.02 | 1.84 |
| 4 | 65.10 | 1.87 | 36.24 | 3.41 | 28.71 | 1.73 | 24.02 | 1.41 |
| 8 | 65.48 | 1.11 | 38.76 | 4.18 | 30.01 | 2.66 | 23.28 | 1.67 |
| 12 | 67.49 | 1.83 | 39.20 | 4.38 | 31.08 | 3.36 | 20.72 | 2.36 |
| 16 | 64.94 | 2.46 | 40.04 | 2.41 | 31.70 | 3.21 | 17.29 | 2.26 |
| 20 | 67.85 | 1.64 | 39.73 | 1.92 | 30.10 | 4.51 | 15.57 | 0.96 |

FRACTIONAL SHORTENING (%)

| FS | SHAM | | CardioCluster | | C+E+M | | Vehicle | |
|---|---|---|---|---|---|---|---|---|
| Week | Mean | SEM | Mean | SEM | Mean | SEM | Mean | SEM |
| Baseline | 34.25 | 2.31 | 35.32 | 1.20 | 35.55 | 1.42 | 34.67 | 1.11 |
| 1 | 38.64 | 1.36 | 14.26 | 2.19 | 14.47 | 1.15 | 13.64 | 1.03 |
| 2 | 36.86 | 1.57 | 15.99 | 1.81 | 14.61 | 1.60 | 12.72 | 1.22 |
| 4 | 34.35 | 1.78 | 16.97 | 2.21 | 12.98 | 0.83 | 11.90 | 0.75 |
| 8 | 36.78 | 1.10 | 17.96 | 2.55 | 13.49 | 1.30 | 10.79 | 1.03 |
| 12 | 37.30 | 1.12 | 18.32 | 3.14 | 15.03 | 1.57 | 10.79 | 0.99 |
| 16 | 35.93 | 2.71 | 18.11 | 1.16 | 14.14 | 1.52 | 8.73 | 1.68 |
| 20 | 40.41 | 2.15 | 19.04 | 1.17 | 14.68 | 2.09 | 8.96 | 1.46 |

**Echocardiographic data represented as mean ± SEM. Heart rate, anterior wall thickness, posterior wall thickness, left ventricular volume, ejection fraction and fractional shortening were measured at specified times after MI. (N) indicates the number of mice used in each group at the given time point.**

**Supplemental Video 1.** CardioCluster spontaneous self-assembly revealed using time-lapse video microscopy.

**Supplemental Video 2.** CardioCluster spontaneous self-assembly with changed seeding sequence so that cCIC+EPCs are added prior to MSC seeding.

#### Supplemental Figure 1. *In vitro* lineage assessment relative to the non-cardiac controls HUVECs and BM MSCs

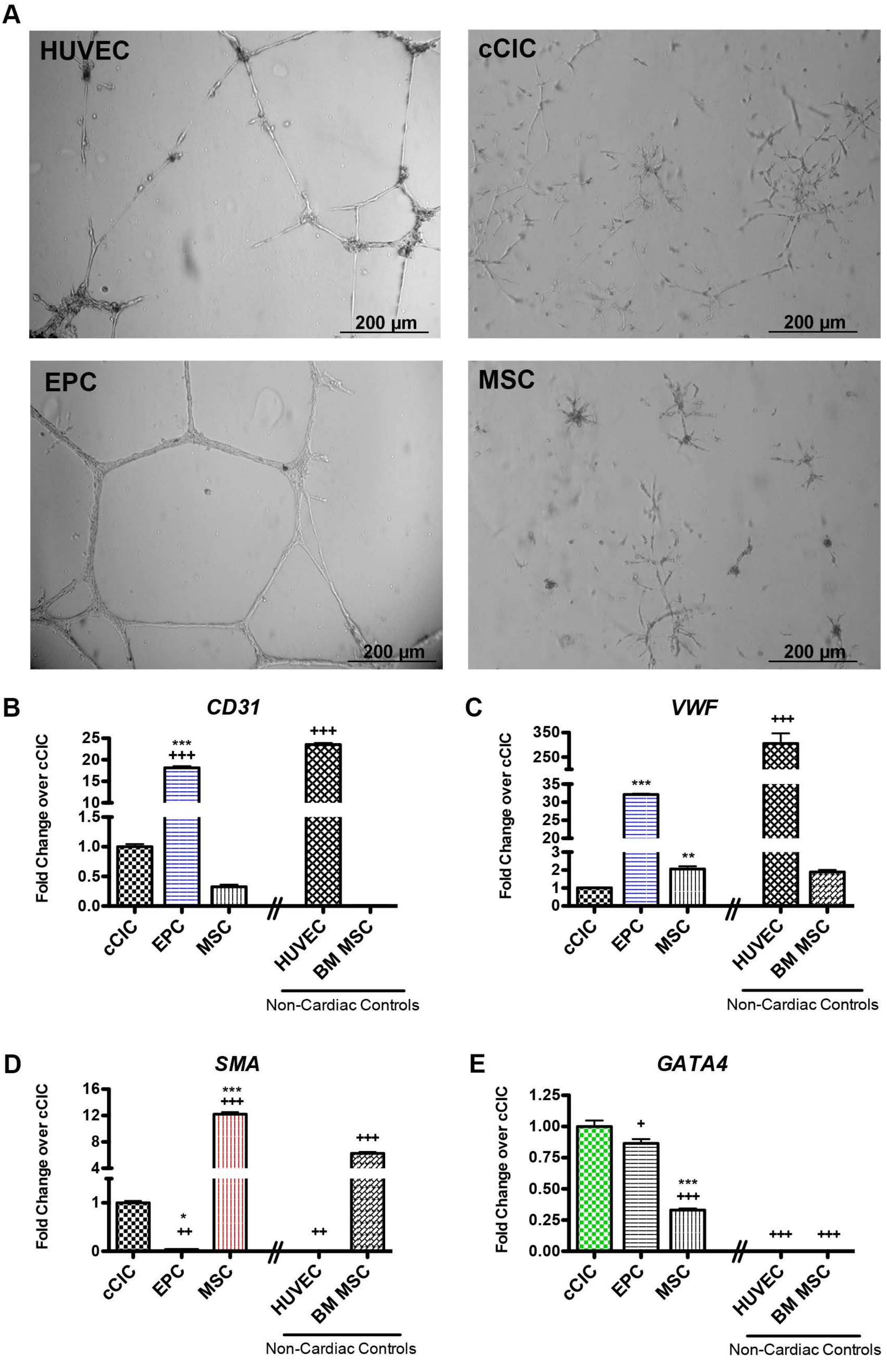

**Supplemental Figure 2.** CardioCluster formation and characterization

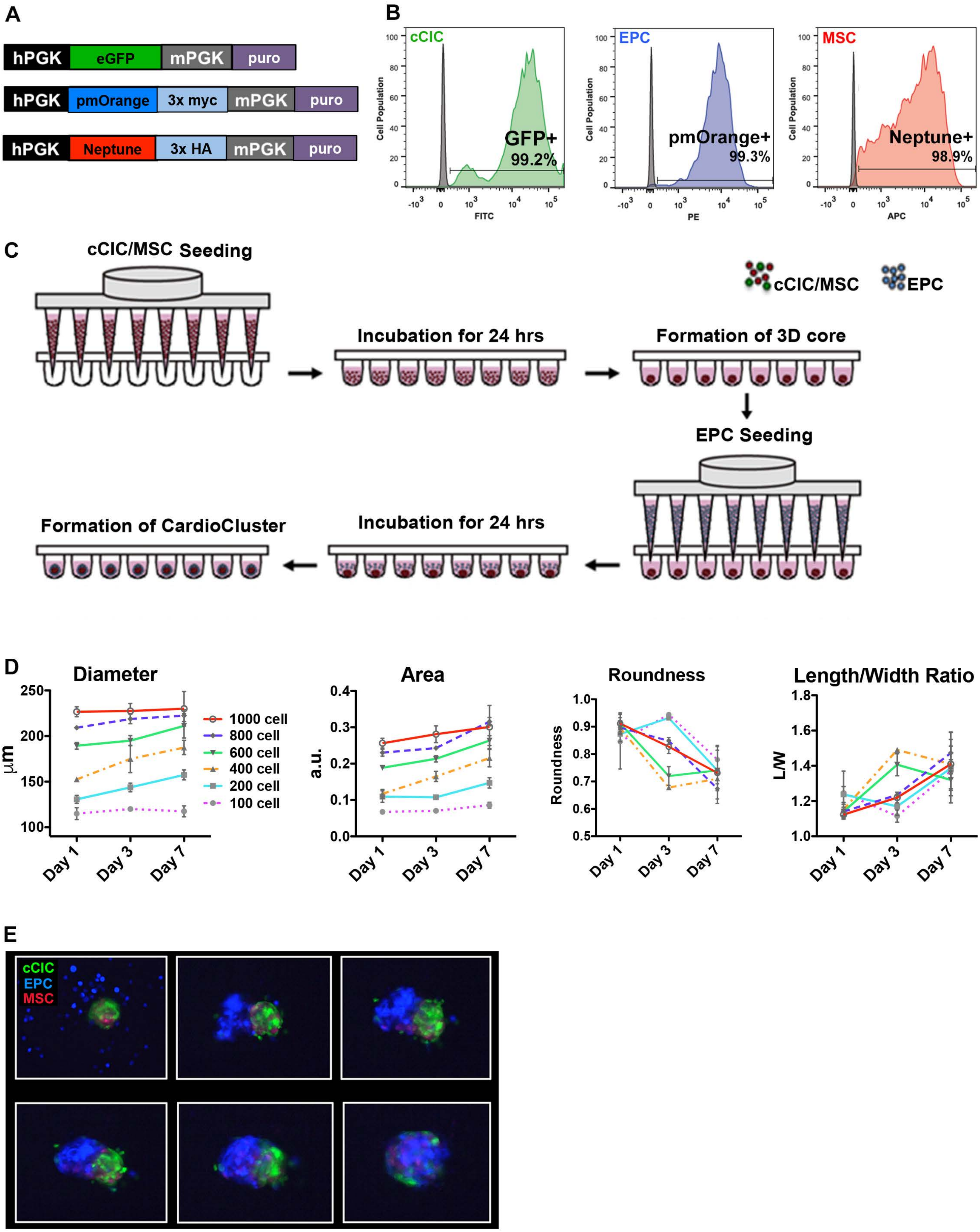

#### Supplemental Figure 3. Cell morphology measurements

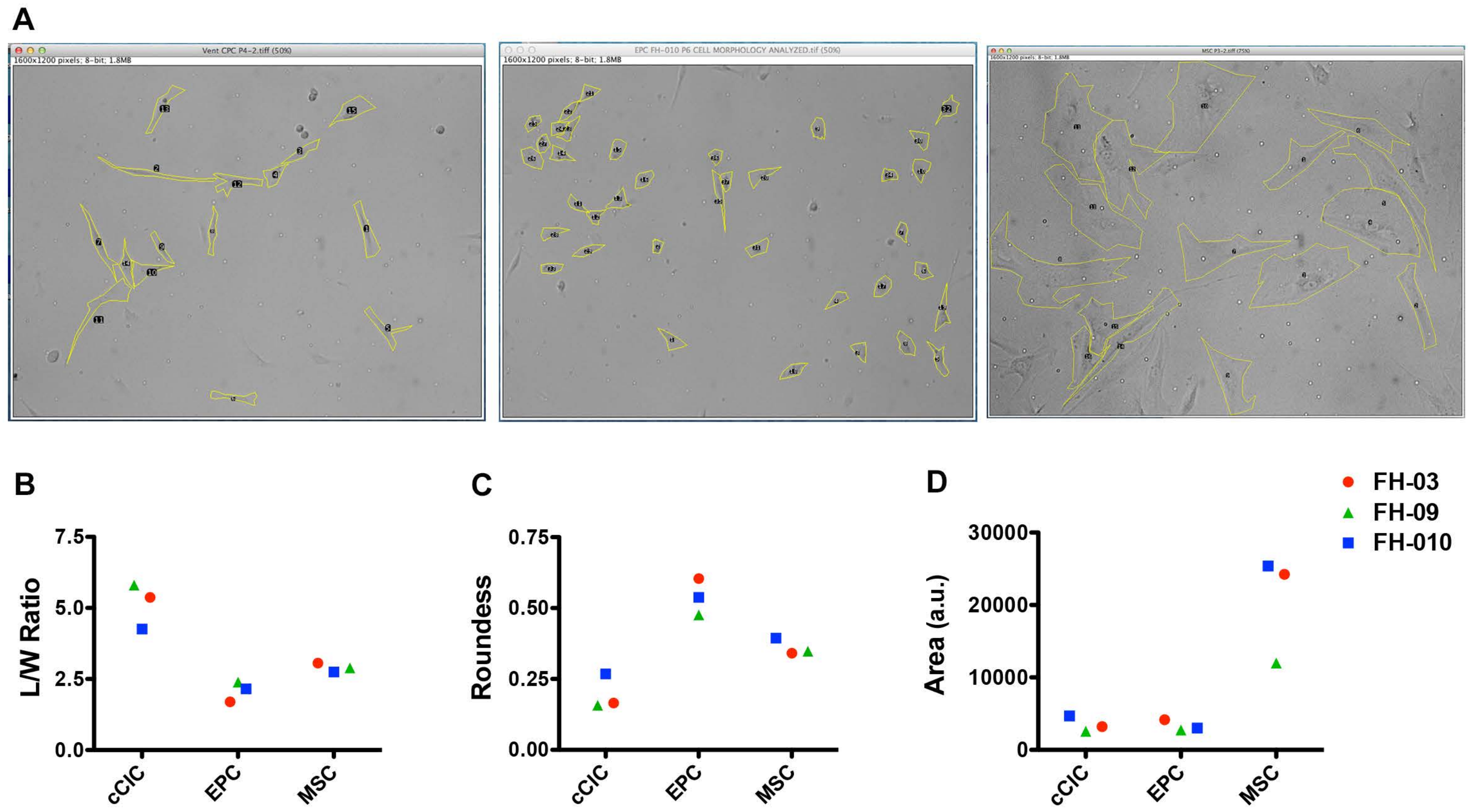

**Supplemental Figure 4. CardioClusters frozen in liquid nitrogen maintain structural integrity and viability**

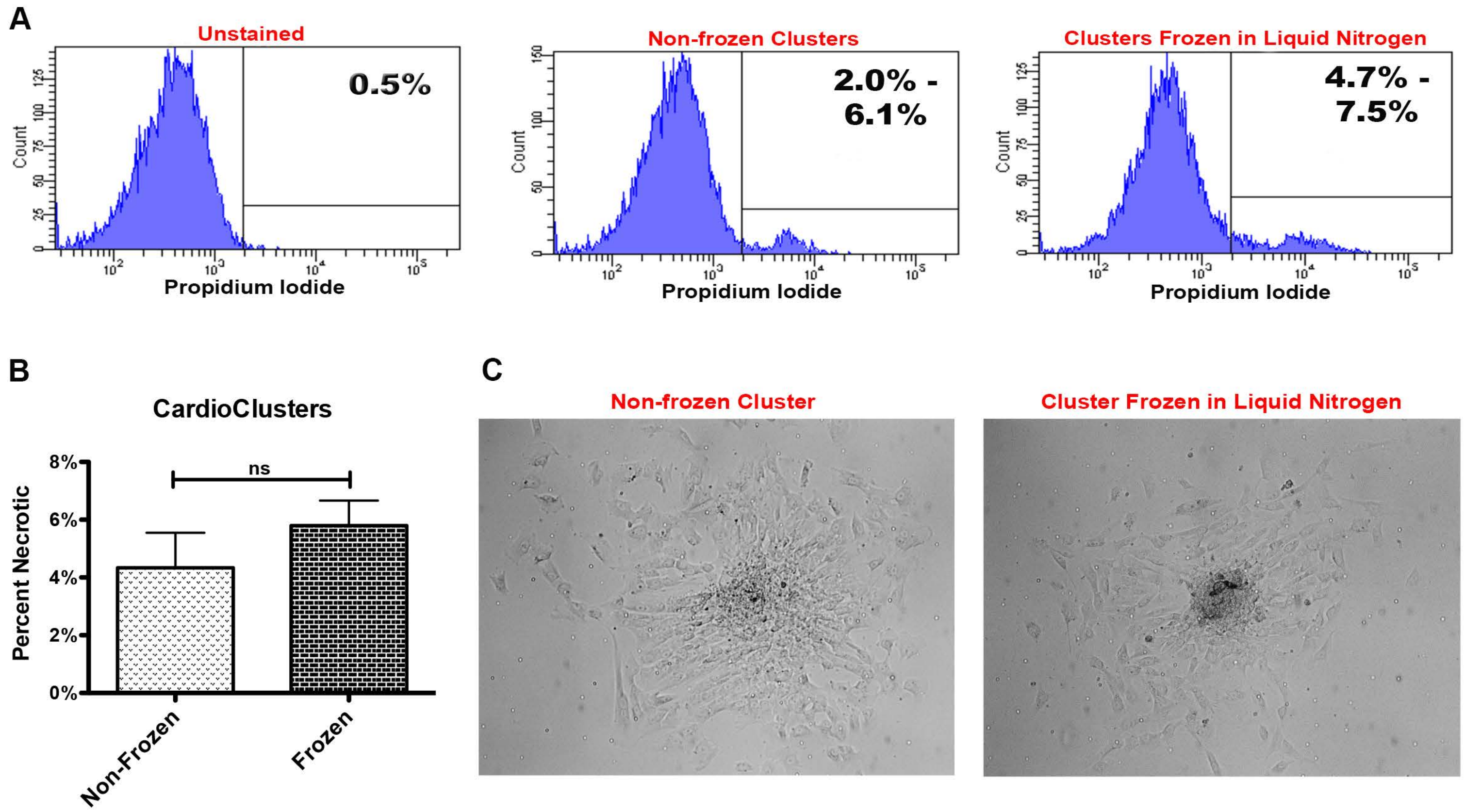

### Supplemental Figure 5. Quality control for single-cell RNA sequencing

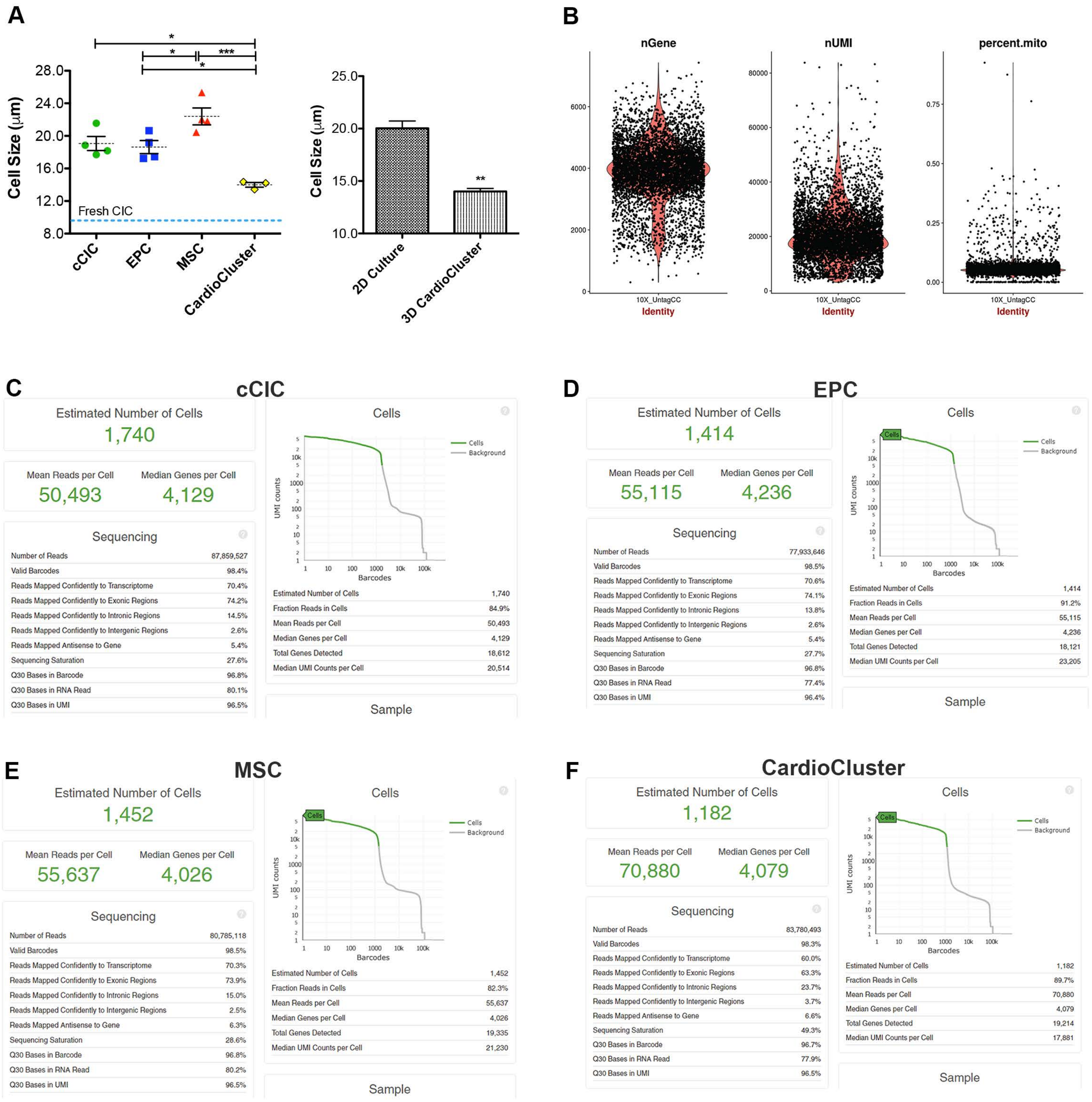

**Supplemental Figure 6.** **CardioClusters restore NRCM morphology following serum starvation**

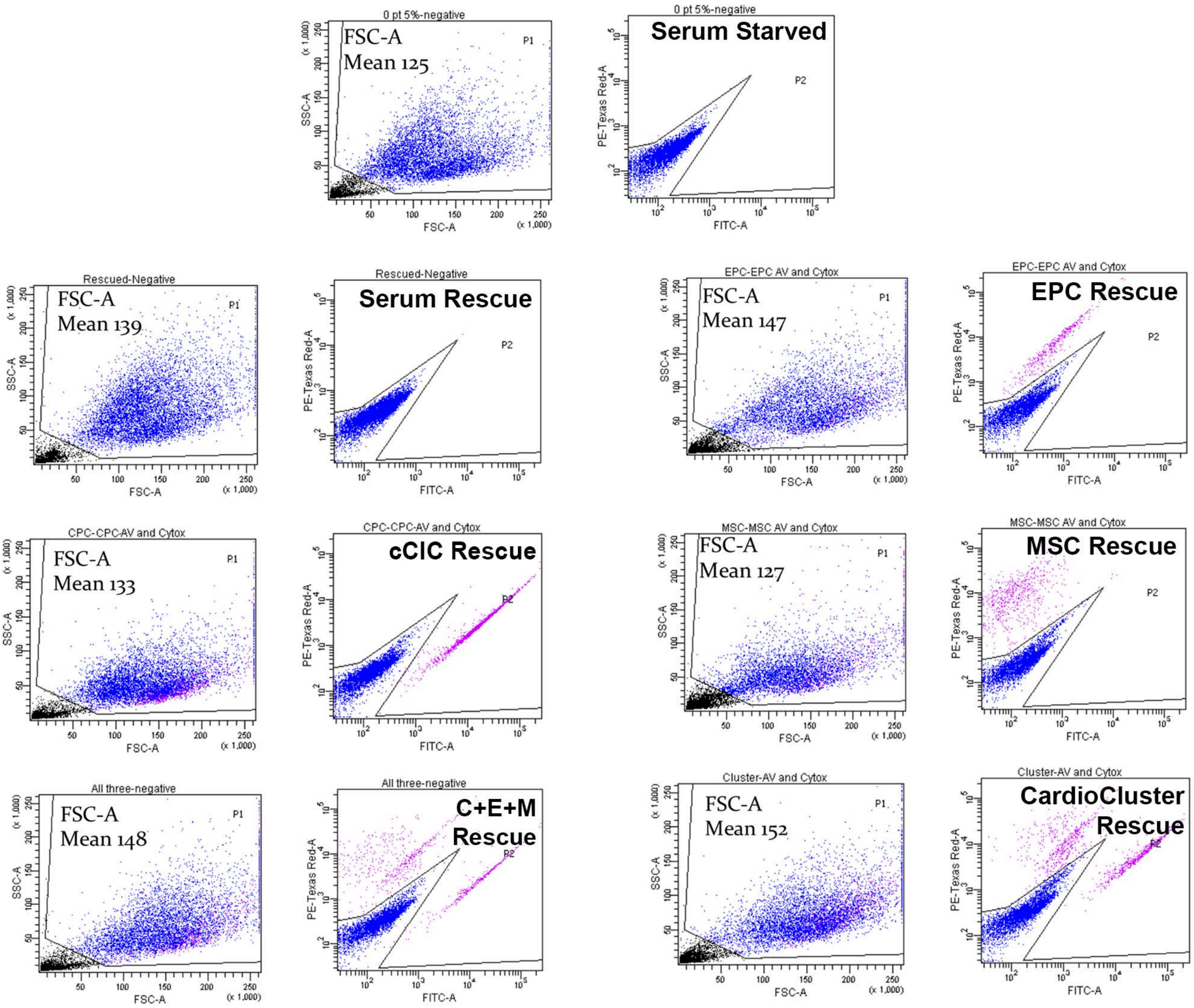

**Supplemental Figure 7.** **CardioClusters show increased paracrine gene expression and remain uncommited towards a particular lineage after *in vitro* co-culture with cardiomyocytes**

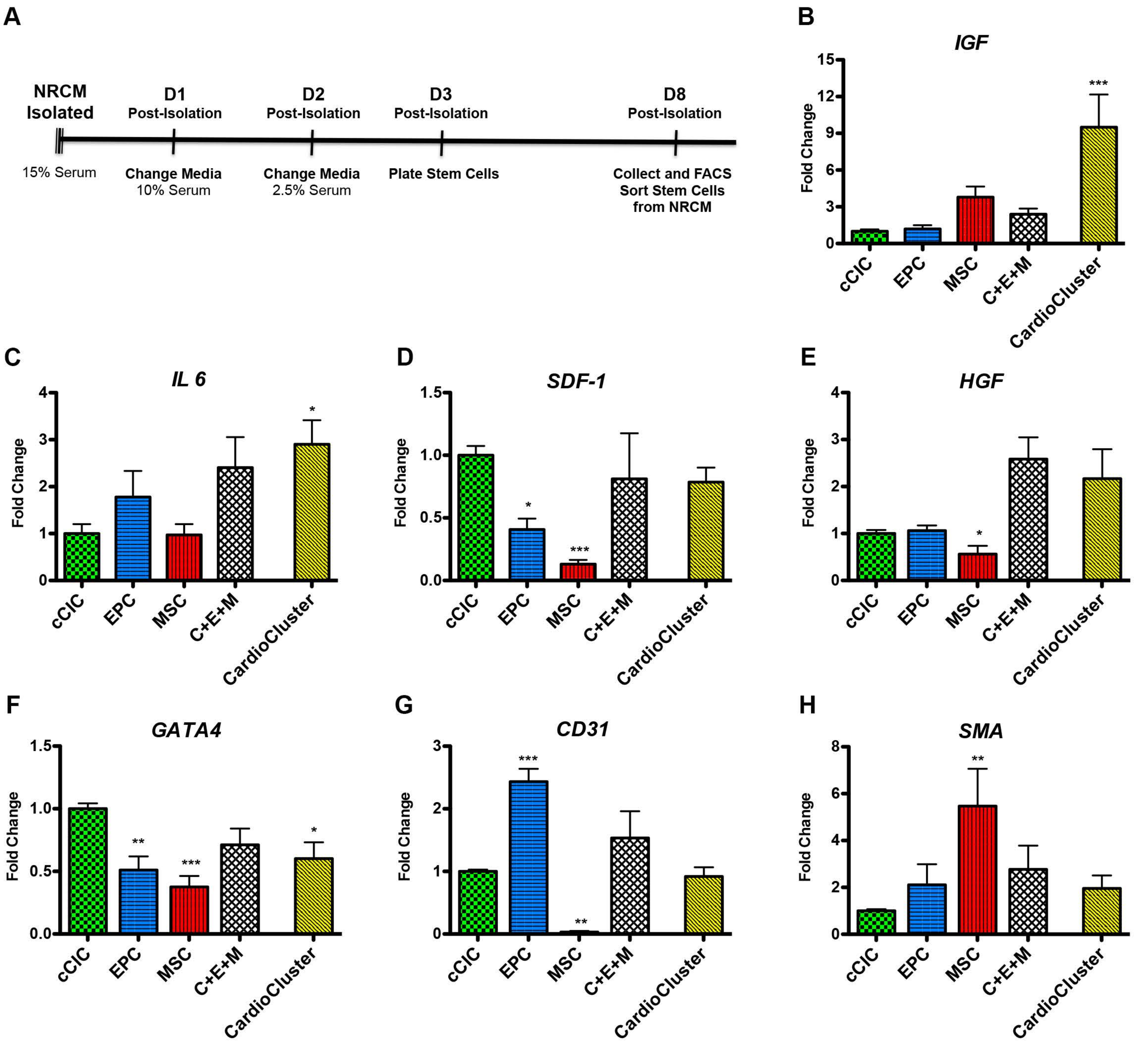

**Supplemental Figure 8.** **Representative staining for markers of apoptosis and necrosis**

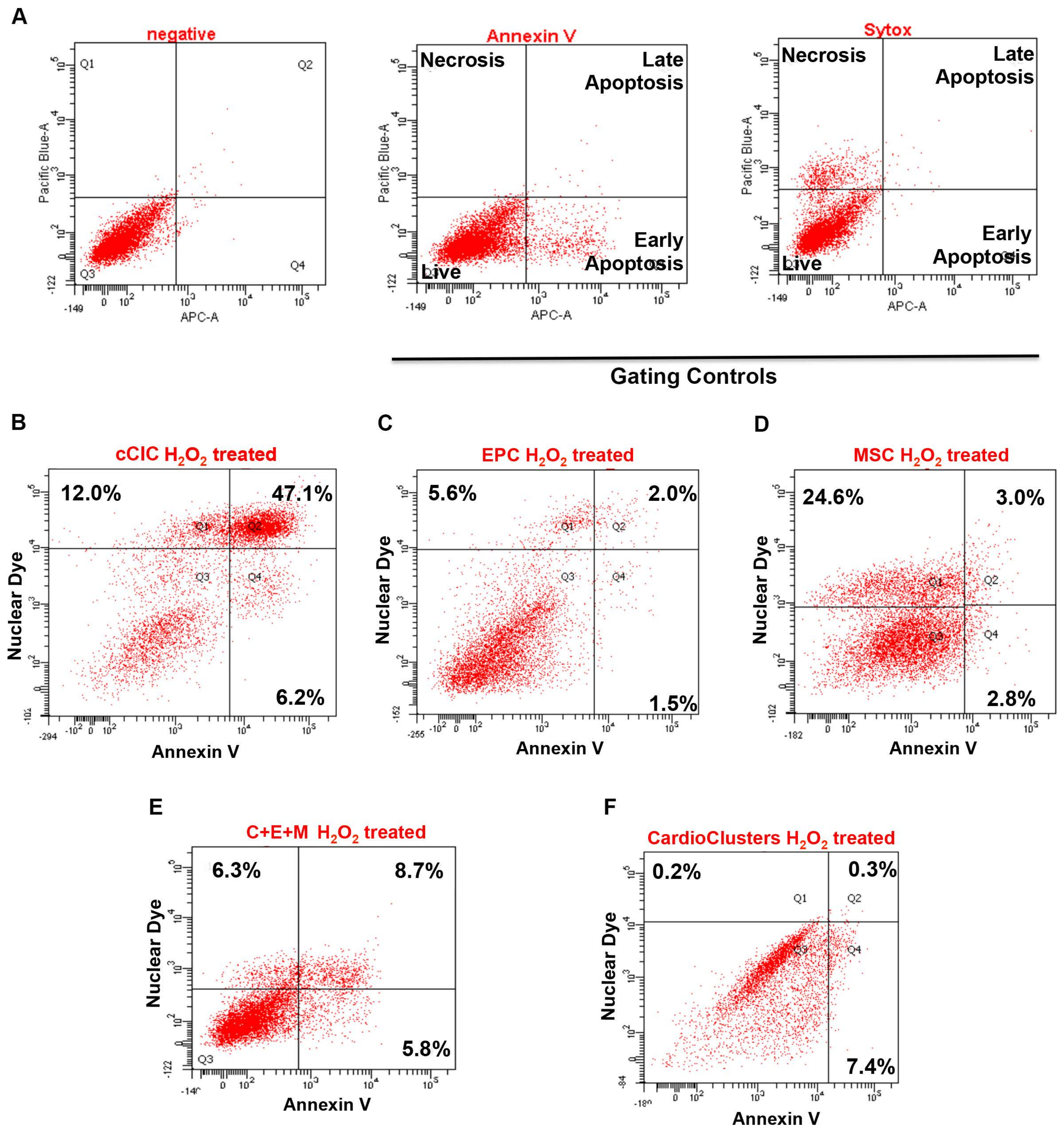

**Supplemental Figure 9.** **Ejection fraction for individual mice grouped by surgery**

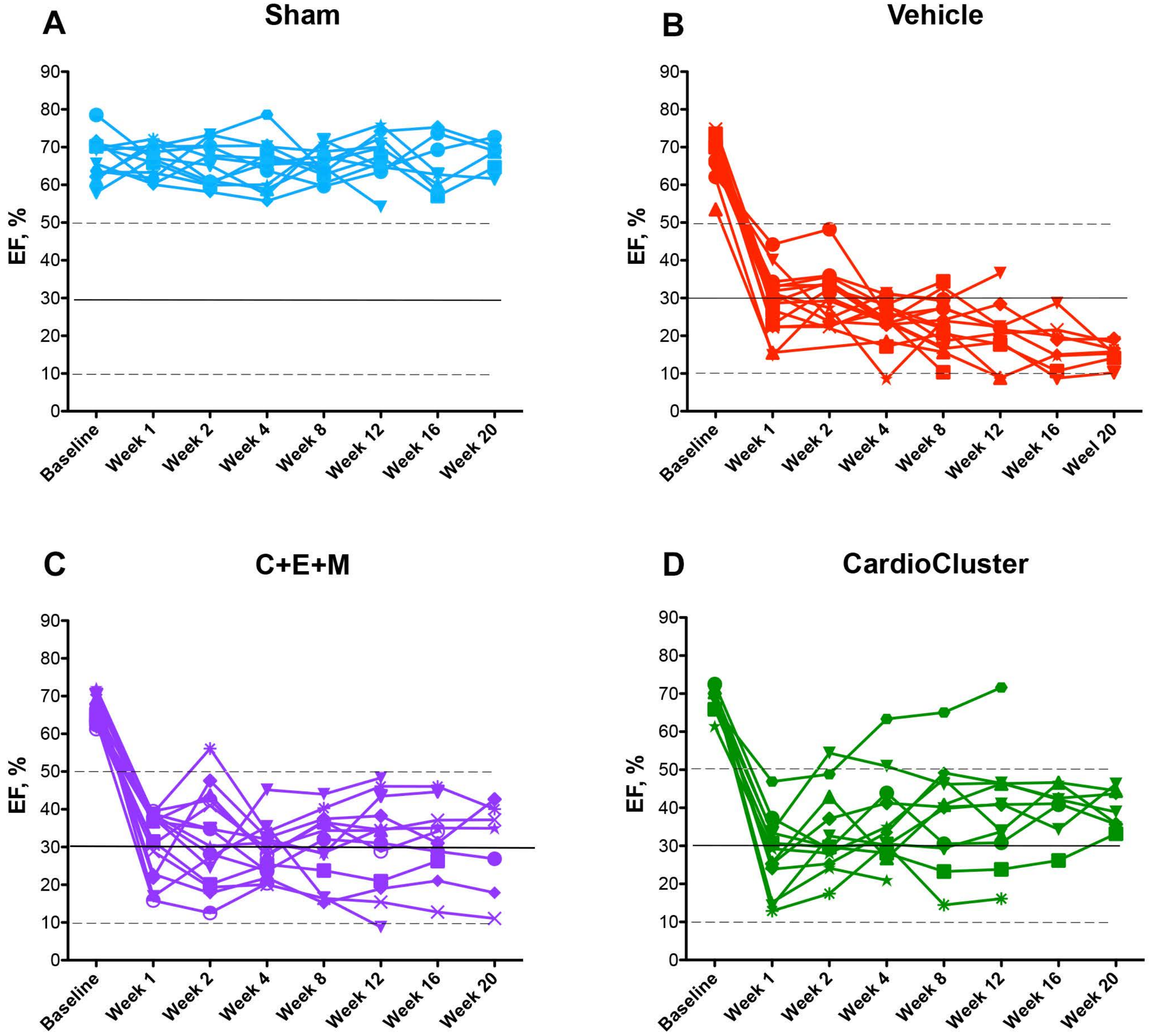

**Supplemental Figure 10.** **Echocardiographic and hemodynamic data confirms CardioCluster treatment improves cardiac function**

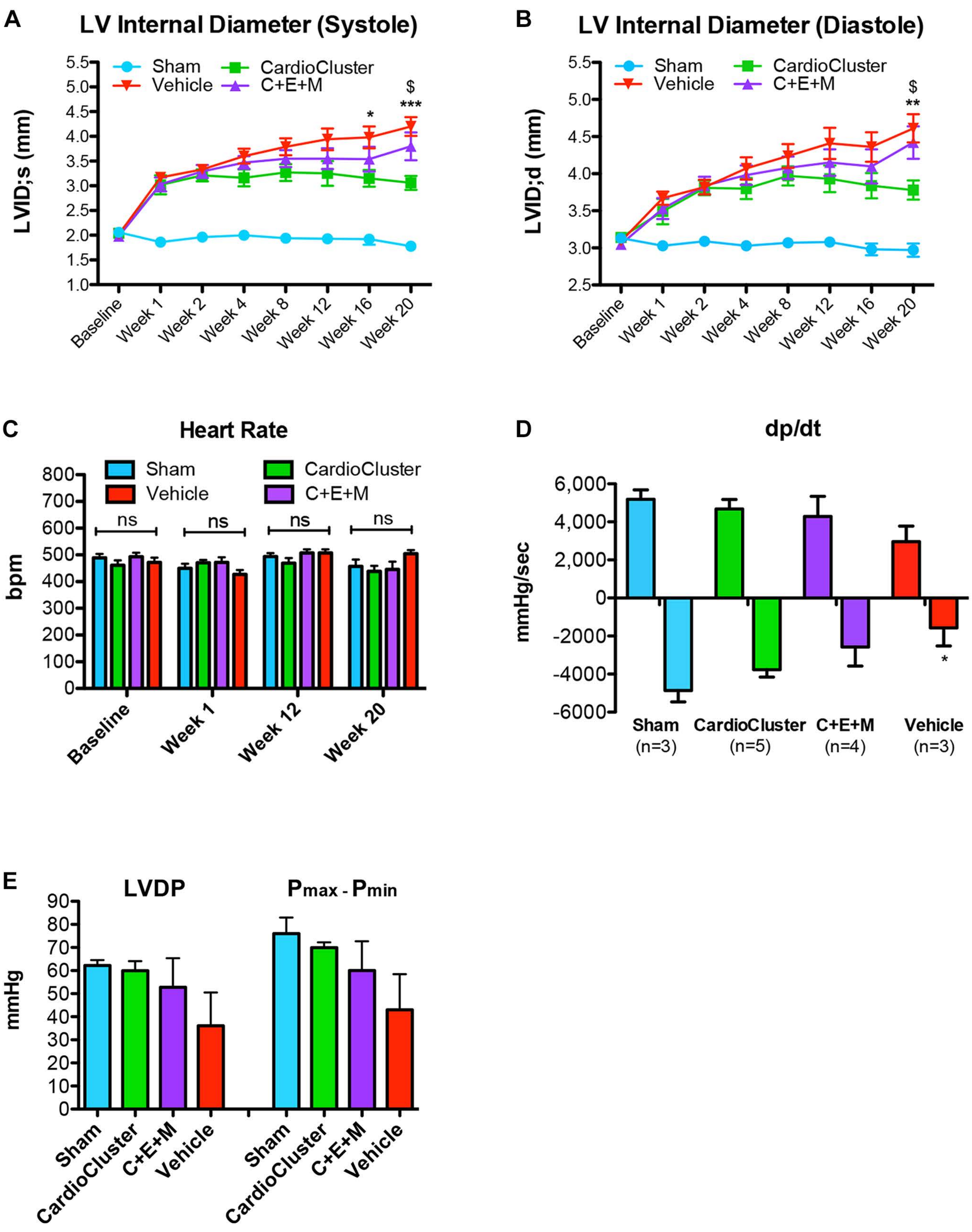

**Supplemental Figure 11.** **CardioCluster treatment antagonizes cardiomyocyte hypertrophy in the border zone and remote region and preserves cardiomyocyte size in the infarcted region**

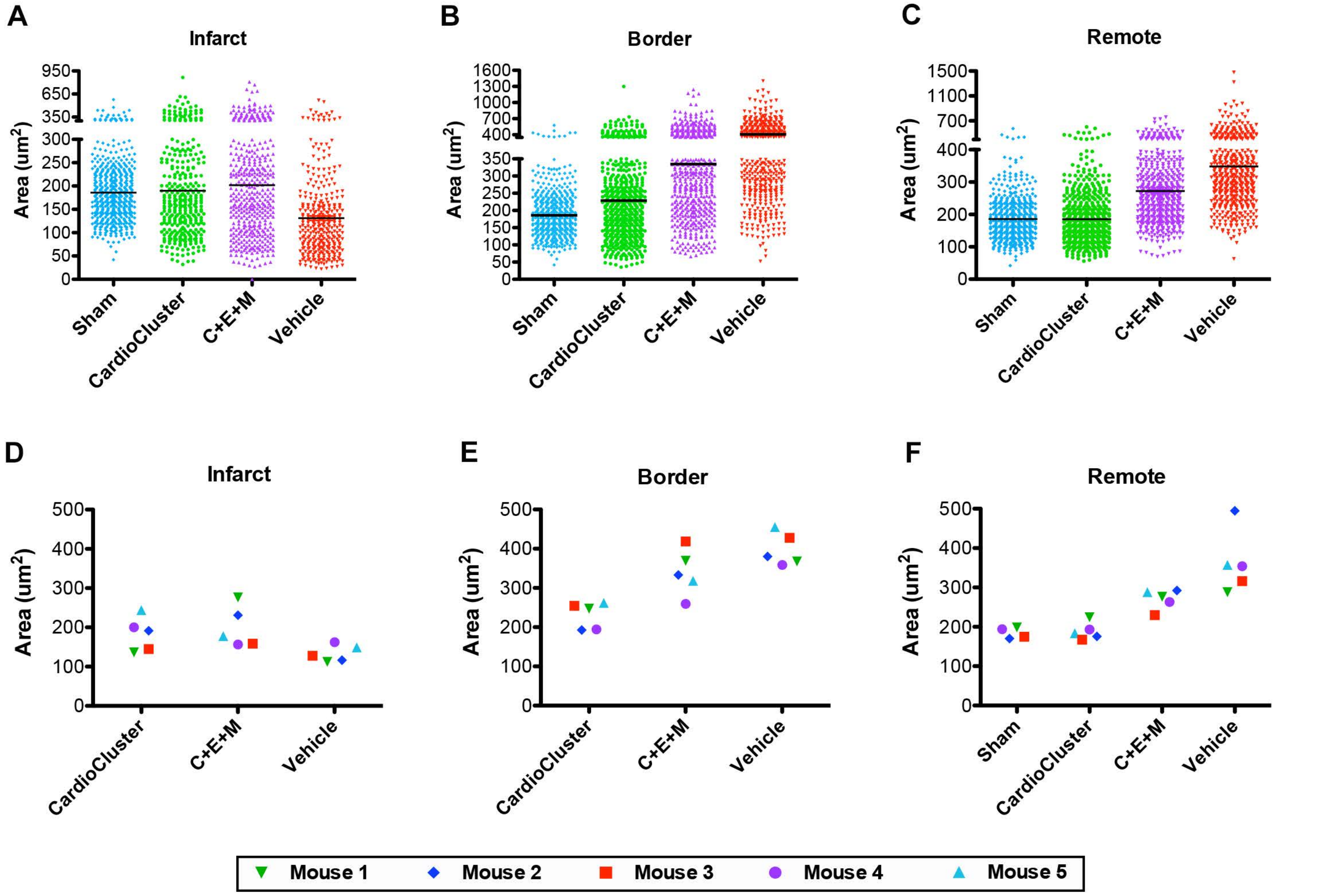
